# Precision Diffusion Imaging

**DOI:** 10.1101/2021.02.19.432023

**Authors:** Nicole A Seider, Babatunde Adeyemo, Ryland Miller, Dillan J Newbold, Jacqueline M Hampton, Kristen M Scheidter, Jerrel Rutlin, Timothy O Laumann, Jarod L Roland, David F Montez, Andrew N Van, Annie Zheng, Scott Marek, Benjamin P Kay, G Larry Bretthorst, Bradley L Schlaggar, Deanna J Greene, Yong Wang, Steven E Petersen, Evan M Gordon, Abraham Z Snyder, Joshua S Shimony, Nico U F Dosenbach

## Abstract

Diffusion tensor imaging (DTI) aims to non-invasively characterize the anatomy and integrity of the brain’s white matter fibers. To establish individual-specific precision approaches for DTI, we defined its reliability and accuracy as a function of data quantity and analysis method, using both simulations and highly sampled individual-specific data (927-1442 diffusion weighted images [DWIs] per individual). DTI methods that allow for crossing fibers (BedpostX [BPX], Q-Ball Imaging [QBI]) estimated excess fibers when insufficient data was present and when the data did not match the model priors. To reduce such overfitting, we developed a novel crossing-fiber diffusion imaging method, Bayesian Multi-tensor Model-selection (BaMM), that is designed for high-quality repeated sampling data sets. BaMM was robust to overfitting, showing high reliability and the relatively best crossing-fiber accuracy with increasing amounts of diffusion data. Thus, the choice of diffusion imaging analysis method is important for the success of individual-specific diffusion imaging. Importantly, for potential clinical applications of individual-specific precision DTI, such as deep brain stimulation (DBS), other forms of neuromodulation or neurosurgical planning, the data quantities required to achieve DTI reliability are lower than for functional MRI measures.

## 1. INTRODUCTION

Brain function is critically dependent on white matter tracts for inter-lobe communication [1]. Studies of white matter tracts connecting distant regions of the brain have greatly advanced our understanding of systems-level brain organization [2]. Damage to white matter via dysmyelination, demyelination, stroke, or trauma, is a key feature of many neurological disorders [3–6].

Diffusion tensor imaging (DTI) is an MRI technique that provides information about water diffusion, which that can in turn be used to probe white matter organization. DTI entails acquisition of multiple diffusion weighted images (DWI), each of which is sensitized to water diffusion in a particular direction. At least six orthogonally oriented DWIs are required to estimate a single diffusion tensor representing the orientation of white matter fibers at a given location in the brain [7–9]. Several shape and orientation characteristics may be extracted from the estimated diffusion tensor: fractional anisotropy (FA), radial diffusivity (RD), axial diffusivity (AD), mean diffusivity (MD), and orientation angles in terms of polar coordinates *θ* and *ϕ*. While a model describing a single tensor is theoretically adequate for simple fiber pathways, the single-tensor model has well recognized limitations for describing the complex, often interleaved orientation of multiple crossing fiber pathways in the human brain. By contrast, more complex models potentially can account for multiple diffusion compartments and thus resolve crossing fibers [10–15].

Early DTI studies acquired the minimum requirement of six orthogonal DWIs for computing a single diffusion tensor [9]. With improvements in MRI hardware and software and the demand for more complex DTI models, acquisition schemes have increased in complexity. Clinical DTI studies typically acquire 12-30 DWIs per patient, while research studies typically acquire 30-60 DWIs per participant [16]. Recent large, multi-site studies such as the Human Connectome Project (HCP, [17]) and the Adolescent Brain Cognitive Development (ABCD, [18]) study, collected 297 and 103 DWIs per participant, respectively. Collecting even more data per individual, through repeated sampling has been informative for functional MRI (precision functional mapping [PFM]) [19–21]. PFM has revealed individual variants in functional network architecture that went undetected with typical amounts of data per subject [22–25] By analogy, intensive acquisition of DWIs in individuals could be similarly fruitful in the study of structural brain connectivity. Earlier studies have examined reliability and accuracy in diffusion imaging studies using less than 60 diffusion directions [26, 27], by comparing mean FA [16, 28, 29], reliability of tract-averaged FA [30, 31], and capacity to resolve crossing-fiber models [32, 33]. However, it is unclear what degree of within-individual reliability could be achieved by collecting much larger quantities of DTI data.

Therefore, we acquired repeated DTI scans over multiple sessions. Three individuals were scanned on multiple days and we acquired 9 - 14 complete DWI datasets per individual using the ABCD study sequence [18]. This sequence includes 103 DWIs (7 B0; 4 b-value shells; ~6.5 minutes). These repeated sampling data were used to study how DTI data quantity and analysis methods impact reliability and accuracy. We pseudo-randomly sampled DWI encodings in a manner that maintained approximately constant angular coverage (see Methods for details, and validation in Figure S1), to systematically evaluate reliability using varying number of spatially distributed DWI. While earlier work had suggested 30 spatially distributed DWIs could be sufficient to estimate a diffusion tensor [27], more complex models had not been similarly tested.

Three crossing-fiber models were compared: Bayesian Multi-Tensor Model-selection (BaMM), the novel method we developed, FSL’s BedpostX (BPX) [10, 11, 34], and Constant Solid Angle Q-ball Imaging (QBI; [12, 35]. As a control, we also tested two single-tensor estimation methods: linear least squares (LLS) and single-tensor Bayesian (STB) [7, 8, 36]. Pertinent model estimation differences may be summarized as follows: BaMM and BPX both use a partial volume models assuming a variable number of radially symmetric fiber compartments. BaMM incorporates a model selection approach to estimate the number of fiber compartments. BPX uses automatic relevance determination (ARD) to down-weight unnecessary fiber compartments. QBI estimates the diffusion orientation distribution function (ODF) in terms of spherical harmonics. We assessed diffusion model estimation accuracy and reliability as a function of data quantity in both real and simulated data.

## 2. METHODS

### 2.1. Voxelwise Parameter Estimation

We evaluated five parameter estimation methods: two methods (Bayesian Multi-Tensor Model-Selection [BaMM] and FSL’s BedpostX [BPX]) used the ball and sticks model [10]; the third crossing fiber method (Constant Solid Angle Q-Ball Imaging [QBI]) used spherical harmonics [35]; and two methods (Linear Least Squares [LLS] and Single Tensor Bayesian [STB]) used the classic single tensor model [7].

#### 2.1.1. Bayesian Multi-Tensor Model-Selection (BaMM) modeling Ball and Sticks

We developed a Bayesian model selection algorithm followed by parameter estimation of the winning model [36]. BaMM evaluated three competing models derived from the ball and sticks model (zero, one, or two fibers in Eq. 1). Model selection and parameter estimation used a Markov-Chain Monte Carlo (MCMC), with Metropolis-Hastings sampling, and simulated annealing procedure. The model selection penalty was scaled based on the input data size.

#### 2.1.2. FSL’s BedpostX [BPX])

The ball and stick model, developed by FSL [10], is an alternative to the single diffusion tensor model [10, 11]. This model is a multi-compartment model, in which the first compartment models the diffusion of free water as isotropic (ball), and the rest of the *k* compartments model diffusion along several axial fiber directions with zero diffusion in the radial direction (sticks). The predicted diffusion signal is:

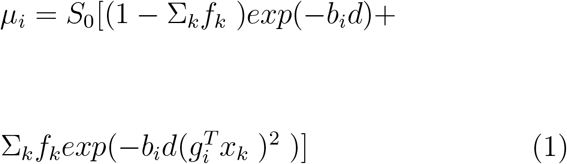

where *i* indexes encoding direction and *k* indexes compartment. *S*_0_ is the signal with no diffusion weighting and *μ_i_* is the signal with a diffusion gradient applied along the unit vector *g_i_* with b-value *b_i_* on diffusion signal *d*. The *f_k_* are volume fractions for each fiber compartment. Each fiber compartment is modeled as a stick-like tensor oriented along *x_k_*. We employed FSL’s BedpostX 6.0.0 to evaluate BPX [34]. The Bayesian parameter estimation approach uses Automatic Relevance Determination (ARD) to down weight unnecessary fibers. BPX estimates angles *θ* and *φ* but not FA, MD, AD, or RD. Angles *θ* and *ϕ* are estimated for every direction (indeed by *k*). We ran BedpostX using the default settings unless noted otherwise: 2 fibers, weight = 1, and burn in = 1000.

#### 2.1.3. Constant Solid Angle Q-Ball Imaging (QBI)

Q-ball imaging is a widely used reconstruction scheme that estimates the orientation distribution function (ODF) through a spherical tomographic inversion [12]. The original Q-ball imaging improved the ODF estimation by considering the constant solid angle (CSA; [35]. E.g., QBI uses spherical deconvolution to estimate the underlying fiber distribution. For ease of comparison to the other methods (LLS, STB, BPX, and BaMM), we estimated the angle of the peaks given by the ODF surface generated by QBI (Figure 1B). Peaks were selecting based on a normalized ODF probability greater than 0.3 and with a matching antipodal peak defined as two peaks having an absolute value dot product greater than 0.99.

**Figure 1:**
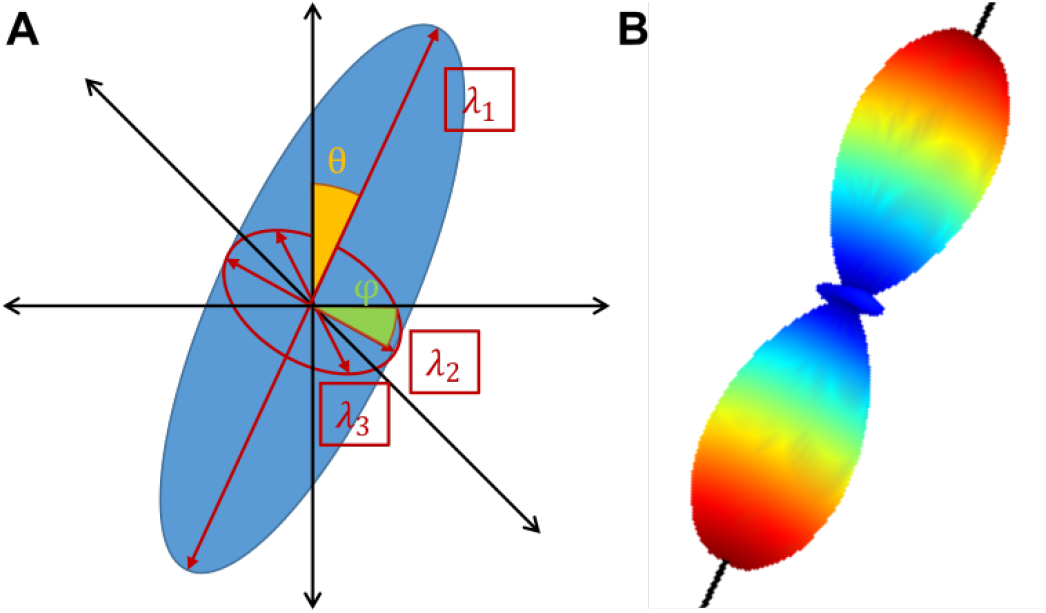
Estimated Tensor and Angles. (A) For LLS and STB, the tensor describing Brownian diffusion of water was calculated. Three eigenvalues are used to describe the tensor shape. From the largest eigenvector, two angles are estimated to describe the tensor orientation in 3D space. For BaMM and BPX, a stick corresponding to eigenvector-1 is estimated and its angles reported. (B) CSA-QBI reports fifteen spherical harmonic values, from which a 3D surface is estimated. The surface is colored by the ODF. The surface/ODF peaks are extracted (black line) and angles *φ* and *θ* estimated to match in A.

#### 2.1.4. Linear Least Squares (LLS)

The LLS method solves an overdetermined system of linear equations by single value decomposition [7, 37]. The solution yields a diffusion tensor *D*, which can be decomposed into eigenvalues (λ_1_, λ_2_, λ_3_, Figure 1A) and eigenvectors (*ν*_1_, *ν*_2_, *ν*_3_). Derived quantities are fractional anisotropy (FA), radial diffusivity (RD), axial diffusivity (AD), mean diffusivity (MD). In addition, the orientation of the principal axis of diffusion can be characterized in terms of polar angles relative to the Z-axis (*θ*) and azimuthal rotation in the XY plane (*ϕ*, Figure 1A [10].

#### 2.1.5. Single Tensor Bayesian Estimation (STB)

The single tensor Bayesian method estimates the posterior probability of the set of parameters, *ω_i_* = (*θ, φ*, λ_1_, λ_2_, λ_3_, *S*_0_) in voxel *j*, given the single tensor model *M* with relevant background information I:

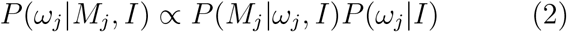

The background information *I* is given as several priors that reflect biological constraints: λ_1_, λ_2_, λ_3_ are limited to between 0 and 3 mm^2^/s, the biological range of diffusion in white matter, and *θ, ϕ* are limited between 0 and *π* owing to the directional symmetry of the diffusion tensor. In STB, the model *M* is the diffusion signal, *μ_i_*, (see below) and tensor, *D*, defined above. To estimate the model parameters, we used standard Monte Carlo Markov Chain methods [36].

### 2.2. Simulated Data

Simulated data were generated using the Gaussian tensor model [7, 8]:

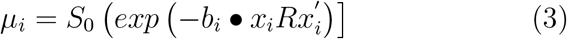

where *S*_0_ is the signal with no diffusion weighting, *μ_i_* is the signal with a diffusion gradient applied along the vector *x_i_* with b-value *b_i_*, and *R* is the diffusion tensor. *S*_0_, was fixed at 1000; *b_i_* and *x_i_* matched twelve acquisitions of the ABCD sequence [18]. Three cases were simulated:

#### 2.2.1. Single Tensor

The first test case simulates highly organized white matter with a single principal direction, as would be expected to be found in the mid-sagittal part of the corpus callosum. *R* was defined to have an anisotropy of 0.86, with angles *θ* and *ϕ* set to 1.8 and 2.8 radians, respectively. Rician noise was added to emulate a signal to noise ratio (SNR) of 30, 50, and 100 [38], to create three data sets at varying SNR.

#### 2.2.2. Two Crossing Tensors

The second test case simulates two highly organized, crossing white matter tracts. The simulations were generated as two highly anisotropic tensors, *R*_1_ and *R*_2_. A range of possibilities was explored by varying the SNR, tensor fraction, FA, and crossing angle of the tensors. Values were varied as follows: SNR = 30, 50, 100; tensor fraction equal weighting, 60%:40%, 70%:30%; FA = 0.6:0.6, 0.6:0.8, 0.8:0.8; crossing angle = 30, 60, 90.

#### 2.2.3. Three Crossing Tensors

The final test case simulates three highly organized, crossing white matter tracts. The simulations were generated as three highly anisotropic tensors, *R*_1_, *R*_2_, and *R*_3_. A range of possibilities was explored by varying the SNR, tensor fraction, FA, and crossing angle of the tensors. Values were varied as follows: SNR = 30, 50, 100; tensor fraction equal weighting, 40%:34%:26%, 53%:34%:13%; FA = 0.6:0.6, 0.7:0.7:0.7, 0.8:0.8:0.8; crossing angle = 30, 60, 90.

### 2.3. Repeatedly Sampled Individual-Specific Data

#### 2.3.1. Participants and Study Design

Three individuals who participated in a study of the effects of arm immobilization functional connectivity contributed data [25]. Participants (25yoF, 27yoM, 35yoM) were scanned daily for two weeks prior to an experimental intervention (unilateral arm casting). Data acquired during and after the casting period are not analyzed in this paper. The Washington University School of Medicine Institutional Review Board provided experimental oversight. Participants provided informed consent for all aspects of the study and were paid for their participation.

#### 2.3.2. MR Image Acquisition

Single-shot echo planar diffusion-weighted MRI was acquired on a Siemens 3T Prisma using a 64-channel head coil. Sequence parameters were: 75 contiguous axial slices, isotropic (2×2×2 mm^3) resolution, TR/TE 3500/83 ms, four shells (b-values 0.25, 0.5, 1.0, and 1.5 ms/mm^2^). This sequence includes 103 volumes and 96 encoding directions. Acquisition time per scan was 6.5 min. Total DWI scans (distributed across scanning sessions) for the three subjects were 9, 12, and 14, resulting in a total of 864, 1140, and 168 total encoding directions, respectively.

#### 2.3.3. DWI Processing

We applied FSL’s Eddy current correction and top-up [39, 40] to each DWI acquisition. Each participant’s DWI was registered to their structural data. During eddy correction, FSL calculated total movement of each DWI relative to the first volume. We excluded volumes with framewise displacement greater than 0.5 mm [41]. The mean and standard deviation of displacement in millimeters relative to the prior volume for each subject were: 0.24 and 0.13 for subject 1; 0.29 and 0.19 for subject 2; and 0.38 and 0.23 for subject 3. Diffusion tensor maps were computed using FSL’s tool DTIFIT [42].

#### 2.3.4. Creation of Reliability Curves Using Permutation Resampling

Model estimation with permutation subsampling was used to quantitatively estimate modeled parameter variability. This approach was used for both simulated data and real human data. All available DWI volumes were concatenated. Subsamples covered the shell surface approximately evenly (Figure 2A). Solid angle sectors were defined by dividing the shell into sixteen bidirectional groups (Figure 2B). The XY-plane was divided into four quadrants and polar angle (*θ*) was divided into four intervals equating arclength. For each permutation, we ensured that all sixteen solid angle sectors were approximately evenly sampled.

**Figure 2:**
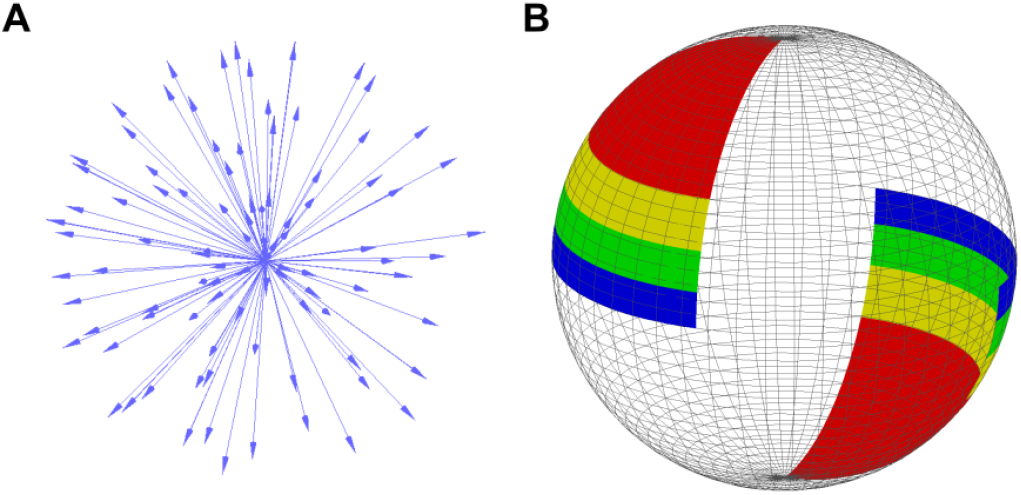
B-Vector selection. Subjects were scanned every day for two weeks, with 96 unique B-vector directions acquired each scan. A) All 1152 B-Vectors from the daily scans plotted on a single sphere. B) B-Vectors were subdivided by their position on the sphere into 16 groups of equal surface area, 4 of which are shown. Encodings were pseudo-randomly selected from the 16 groups to obtain approximately uniform angular sampling over the sphere.

For all parameter estimation methods (BaMM, BPX, QBI, LLS, STB), we compared the estimation of modeled diffusion parameters (*FA, AD, RD, AD, MD, θ, ϕ*} over the range N = 10 up to all available data, in approximately logarithmically spaced increments. DWI volumes were quasi-randomly selected according to the above-described scheme. These steps were repeated over 1000 permutations at each subsampling size. For single tensor shape diffusion parameters (*FA, AD, RD, MD*}, the parameter estimate variability was defined as

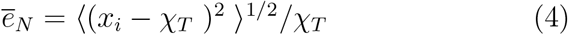

Where *x_i_* represents a parameter estimate (*FA, AD, RD, AD, MD*} obtained from an individual permutation, *χ_T_* is the “true” value obtained using all available data or the ground truth in simulations, and the bracket denotes mean over permutations. *ē_N_* was plotted as a function of sample size (*N*), creating reliability curves for each parameter. Since diffusion is estimated as a bipolar tensor that is symmetric around the origin, the error estimation for angles *θ, ϕ* was modified accordingly to account for modulus pi.

#### 2.3.5. Mean Error Threshold Whole Brain Maps

To generate a voxel-wise heatmap visualizing the threshold sample size *N* needed to reach a mean error less than 5% for each voxel, we conducted the permutation testing described above on every voxel of the brain using the LLS method. The mean error was calculated for each voxel at each value of *N*. A heatmap was created for each diffusion metric, such that voxels are colored by the number of measurements needed to reach a mean error < 5%, where a lower value of N is shown in cool colors, and a larger *N* is in warm colors. If the voxel never reached a mean error < 5% at a sample size of 1000, the voxel is colored in red.

## 3. RESULTS

### 3.1. Single Tensor Simulations: BaMM Detects a Single Fiber More Accurately Than BPX or QBI

Brain anatomy shows that some regions of the brain, such as the corpus callosum, have a single dominant fiber direction. Thus, we first evaluated the DTI methods with simulated single tensor fiber data. To test the accuracy and reliability of diffusion metrics, we used permutation subsampling of the simulated diffusion data to estimate parameter variability for all crossing fiber methods (BaMM [Bayesian Multi-tensor Model-selection], BPX [BedpostX], QBI [Q-Ball Imaging]) and single tensor methods (LLS [Linear Least Squares], STB [Single Tensor Bayesian]). We plotted the estimated radian value of the angles (*ϕ, θ*) to highlight the number of fibers estimated at each DWI sample size. Open circles represent the results of individual permutations and are colored according to the number of fibers estimated (Figure 3).

**Figure 3:**
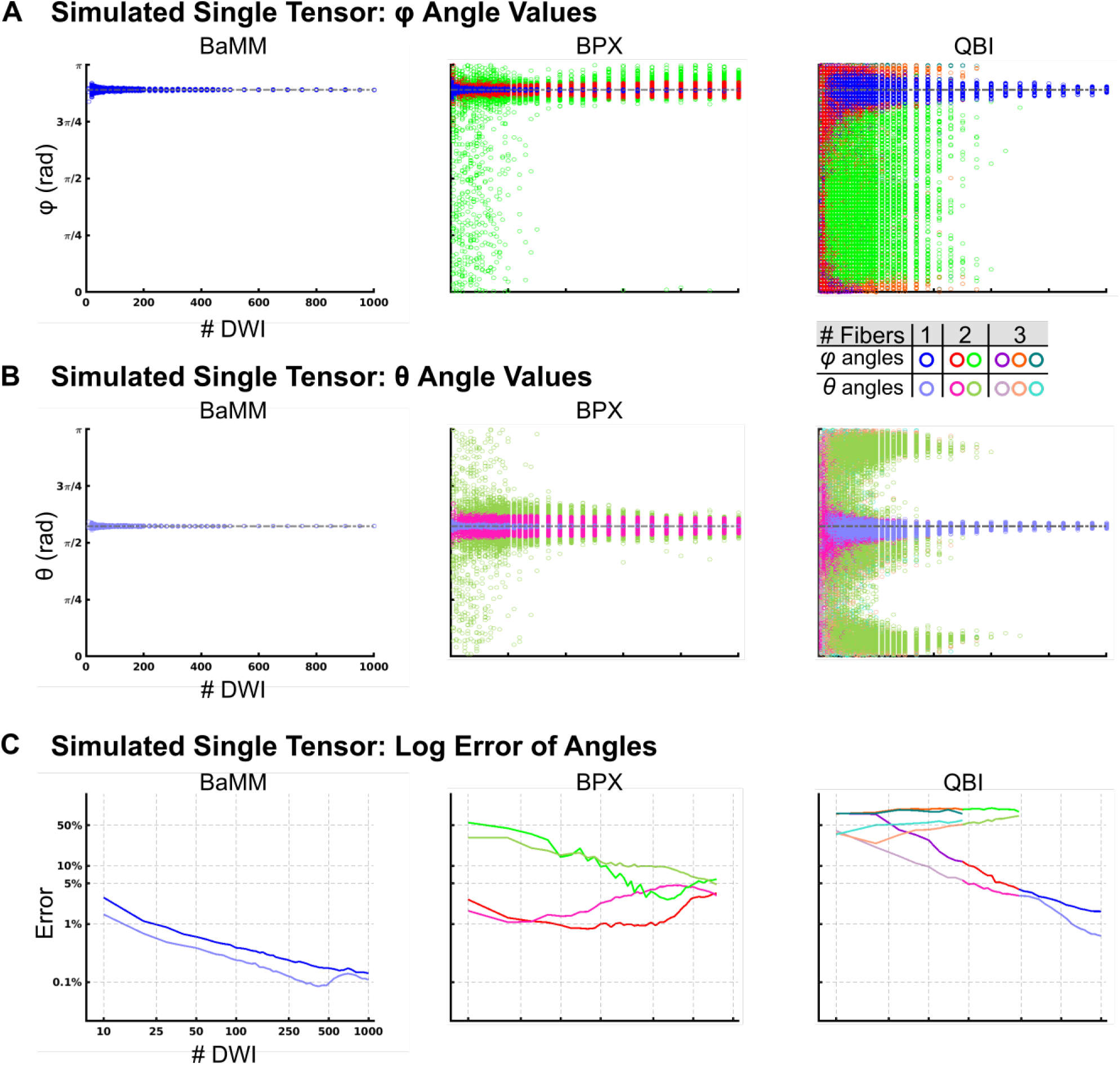
Accuracy of DTI Measures: Simulated Single Tensor. (A) *φ* angle estimations by Bayesian Multi-tensor Model-selection (BaMM), BedpostX (BPX), and constant solid angle Q-Ball Imaging (QBI). Open circles represent the results obtained by repeated permutation sampling. Same color legend for all data panels. Permutations that resulted in a single fiber direction are plotted in blue (*φ*). Permutations that resulted in two fibers are plotted in red (*φ*) and green (*φ*). Permutations that resulted in three fibers are plotted in purple (*φ*), orange (*φ*), teal (*φ*). (B) θ angle estimations. Permutations are plotted in sky blue (*θ*) for one fiber, pink (*θ*) and olive (*θ*) for two fibers, and lilac (*θ*), salmon (*θ*), cyan (*θ*) for three fibers. (C) Error estimation for BaMM, BPX, and QBI. Mean error at each subsampling size was calculated, then plotted on a log scale. The same colors as in (A/B) are used and indicate the most frequent number of fibers estimated at each subsampling size.

Multiple SNR values were tested to track SNR’s effect on reliability and accuracy. The forward-modeled parameter space for simulated single tensors was: SNR 50 (Figure 3), 30, and 100 (Figure S2 for BaMM, BPX, and QBI, and S3 for LLS and STB). Initially, default settings were used for all modeling schemes: BaMM and QBI up to three fibers; BPX two fibers (for same analyses using BPX with other settings see Figure S4).

BaMM accurately estimated the orientation of the single forward modeled principal eigenvector, even with limited quantities of data (> 20 DWI samples; blue/sky blue symbols Figure 3A-B, S2). However, BPX generally falsely estimated two fibers (68.8% of permutations at DWI = 10, at least 90% of permutations at DWI > 120), even when given large quantities of data (Figure 3A-B, S2). At DWI < 200, the angle of the second fiber was broadly distributed over the interval 0 to π (green/olive symbols Figure 3A-B). At DWI > 400, BPX continued to estimate two fibers separated by a small angle, the mean of which closely approximated the single modeled principal eigenvector. When the max number of fibers was increased to 3 (default 2), BPX falsely estimated three fibers in the majority of permutations (39.0% of permutations at DWI = 10, linearly increasing to 88.4% of permutations at DWI = 1000; Figure S4).

QBI estimated one, two or three fibers given different numbers of DWI. At < 90 DWI, QBI was most likely to estimate three fibers that were broadly distributed over the interval 0 to π, and also frequently estimated two or one or two fibers (at 10 DWI, 90% of permutations estimated three fibers, 10% estimated two. By 80 DWI, 47%, 42%, and 11% of permutations estimated three, two, and one fiber respectively). Unlike BPX, QBI consistently and accurately estimated a single fiber at higher DWI quantities (300 DWI: 12%, 31%, and 57% of permutations estimated three, two, and one fiber, respectively. Over 90% of permutations estimated a single fiber at > 460 DWI.

Mean measurement error was calculated relative to the forward-modeled angle or shape metric, to quantify the accuracy of each method as a function of the number of diffusion measurements (Figure 3C, Eq. 4). Log error is plotted by color according to the most frequently estimated number of fibers at each subsampling size. For BaMM, (Figure 3C), error linearly decreased with increasing subsampling size. In contrast, for BPX a linear decrease of error with increasing subsampling measurements was only detected for the secondary fiber (green/olive) but not the primary fiber (red/pink). QBI’s error decreased with increasing subsampling sizes only for the primary fiber, while the second and third fiber had very high errors.

We also evaluated the accuracy of the single tensor methods LLS and STB on simulated single tensor data. As expected and similar to BaMM, LLS and STB estimation of FA, AD, RD, MD, and angles *φ* and *θ* improved with increasing number of diffusion measurements (Figure S3).

### 3.2. Two Tensor Simulations: BaMM Is Overfitting Robust

Next, we simulated two crossing fibers, as commonly detected in deep white matter, e.g., within the crossing of the superior longitudinal fasciculus and uncinate (Figures 4, S5–8). We explored the following forward-modeled parameter space: fiber crossing angle (30°, 60°, 90°), relative weight of fiber compartments (50/50, 60/40, 70/30), SNR (30, 50, 100), and FA of tensors (0.6/0.6, 0.6/0.8, 0.8/0.8). The parameter space was chosen to explore fiber orientation, relative size of fiber compartments, SNR, and the respective FA of the fiber compartments. Figure 4 shows the results of 90° crossing angle, 60/40 relative weight, SNR 50, and FA of 0.8/0.8. Results corresponding to the full parameter space are reported in the Supplemental Figures (Figures S5–7, BaMM, BPX, and QBI respectively). Results were consistent across the parameter space, with slight variations in subsampling size needed to reach specific error thresholds. The single tensor models tested (LLS and STB), estimated the two crossing fibers as a weighted average and the single tensor’s principal eigenvector reflects this inaccurate shape assumption (Figure S8). Again, default settings were used for all modeling schemes: BaMM and QBI up to three fibers; BPX two fibers (for same analyses of BPX using other settings see Figure S9).

**Figure 4:**
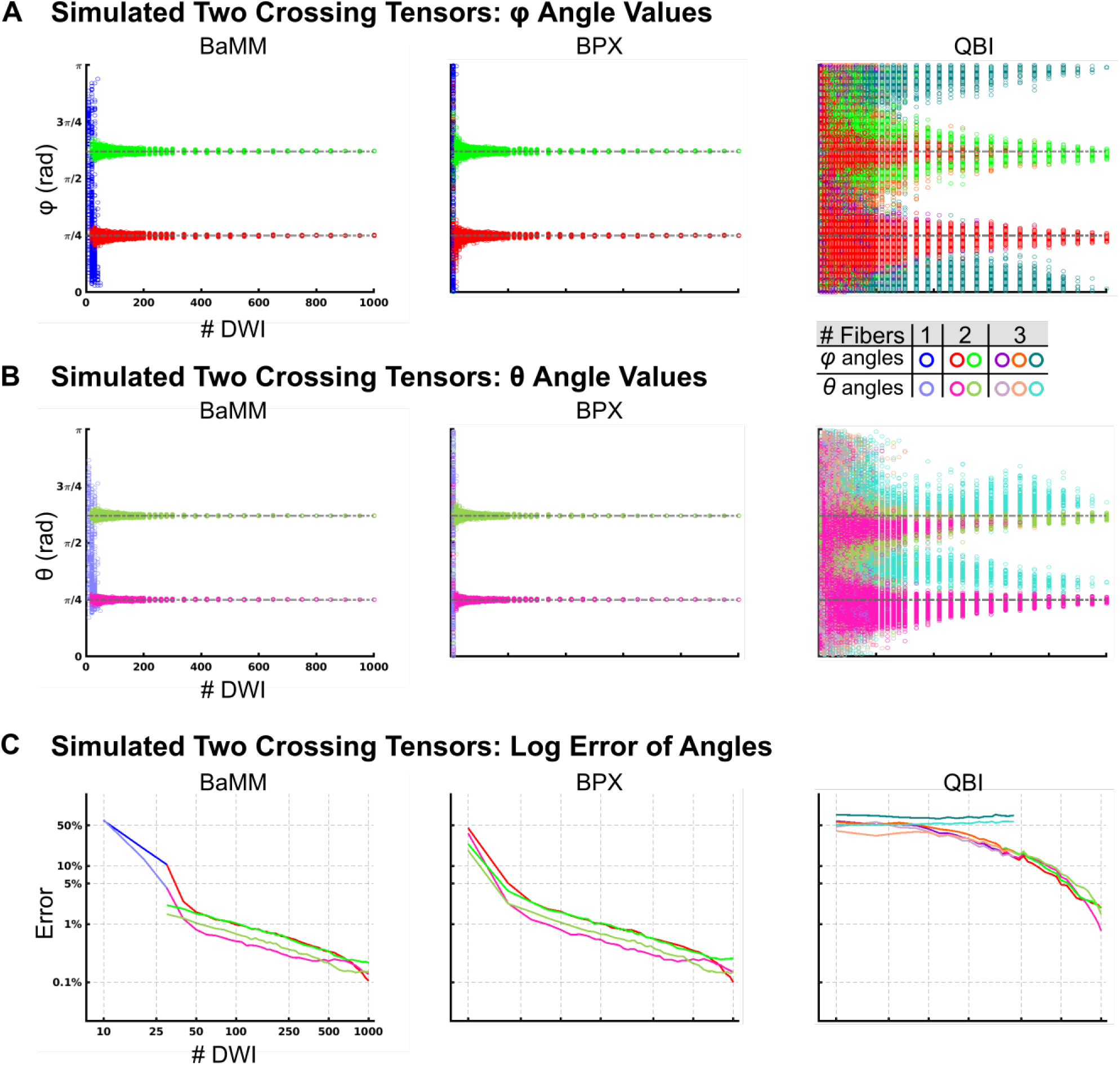
Accuracy of DTI Measures: Simulated Two Crossing Tensors. The tensors were oriented such that they were perpendicular to each other. The first tensor had larger weighting equal to 60% of the signal. Rician noise was added for an SNR = 50. (A) *φ* angle estimations by Bayesian Multi-tensor Model-selection (BaMM), BedpostX (BPX), and constant solid angle Q-Ball Imaging (QBI). Open circles represent the results obtained by repeated permutation sampling. Same color legend for all data panels. Permutations that resulted in a single fiber direction are plotted in blue (*φ*). Permutations that resulted in two fibers are plotted in red (*φ*) and green (*φ*). Permutations that resulted in three fibers are plotted in purple (*φ*), orange (*φ*), teal (*φ*). (B) *θ* angle estimations. Permutations are plotted in sky blue (*θ*) for one fiber, pink (*θ*) and olive (*θ*) for two fibers, and lilac (*θ*), salmon (*θ*), cyan (*θ*) for three fibers. (C) Error estimation for BaMM, BPX, and QBI. Mean error at each subsampling size was calculated, then plotted on a log scale. The same colors as in (A/B) are used and indicate the most frequent number of fibers estimated at each subsampling size.

**Figure 5:**
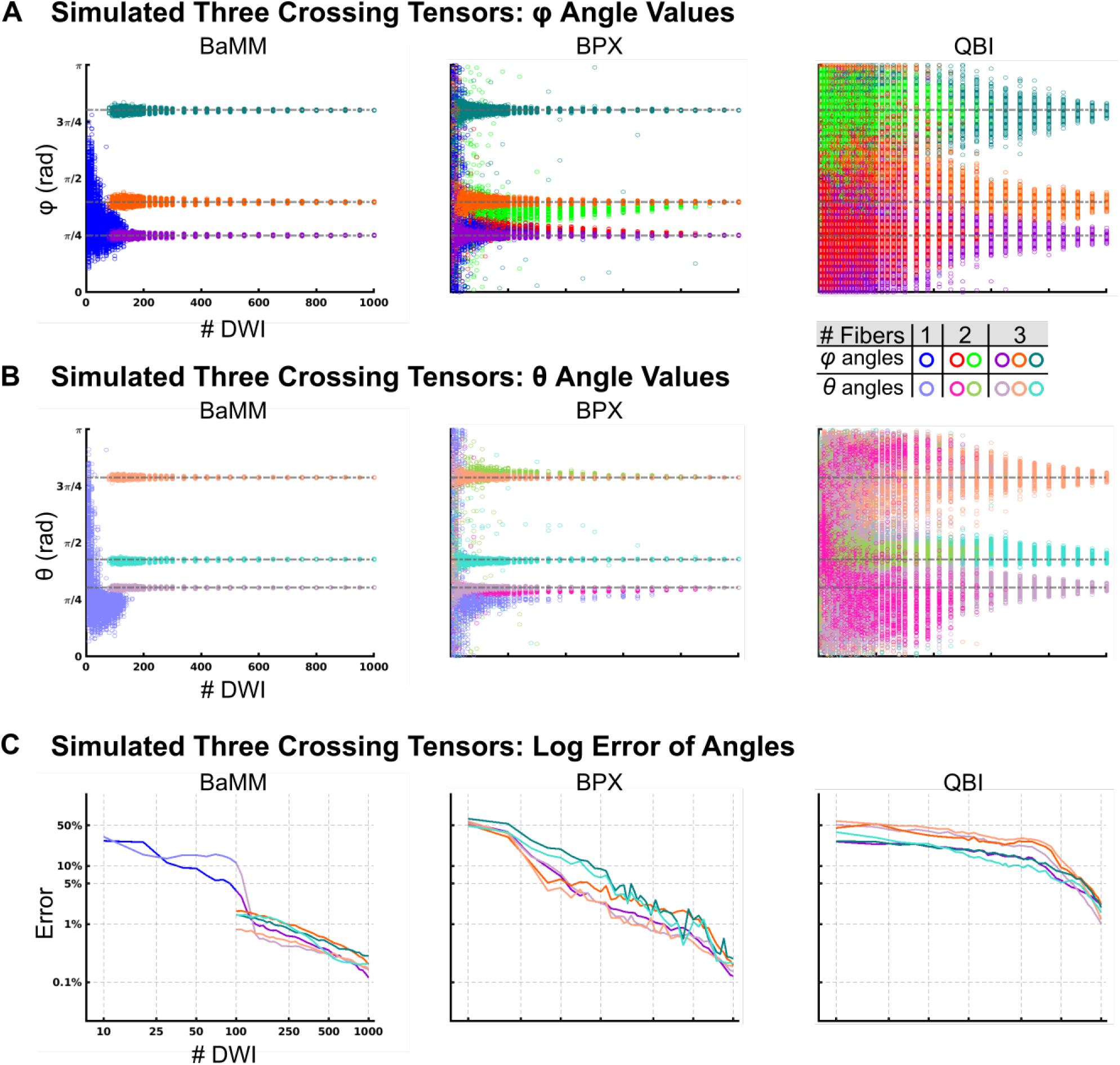
Accuracy of DTI Measures in Simulated Three Crossing Tensor Data. The tensors were oriented such that they were perpendicular to each other. The first tensor had larger weighting equal to 40% of the signal, the second tensor was 34%, and the smallest tensor was 26%. Rician noise was added for an SNR = 50. (A) φ angle estimations by Bayesian Multi-tensor Model-selection (BaMM), BedpostX (BPX), and constant solid angle Q-Ball Imaging (QBI). Open circles represent the results obtained by repeated permutation sampling. Same color legend for all data panels. Permutations that resulted in a single fiber direction are plotted in blue (φ). Permutations that resulted in two fibers are plotted in red (φ) and green (φ). Permutations that resulted in three fibers are plotted in purple (φ), orange (φ), teal (φ). (B) θ angle estimations. Permutations are plotted in sky blue (θ) for one fiber, pink (θ) and olive (θ) for two fibers, and lilac (θ), salmon (θ), cyan (θ) for three fibers. (C) Error estimation for BaMM, BPX, and QBI. Mean error at each subsampling size was calculated, then plotted on a log scale. The same colors as in (A/B) are used and indicate the most frequent number of fibers estimated at each subsampling size

**Figure 6:**
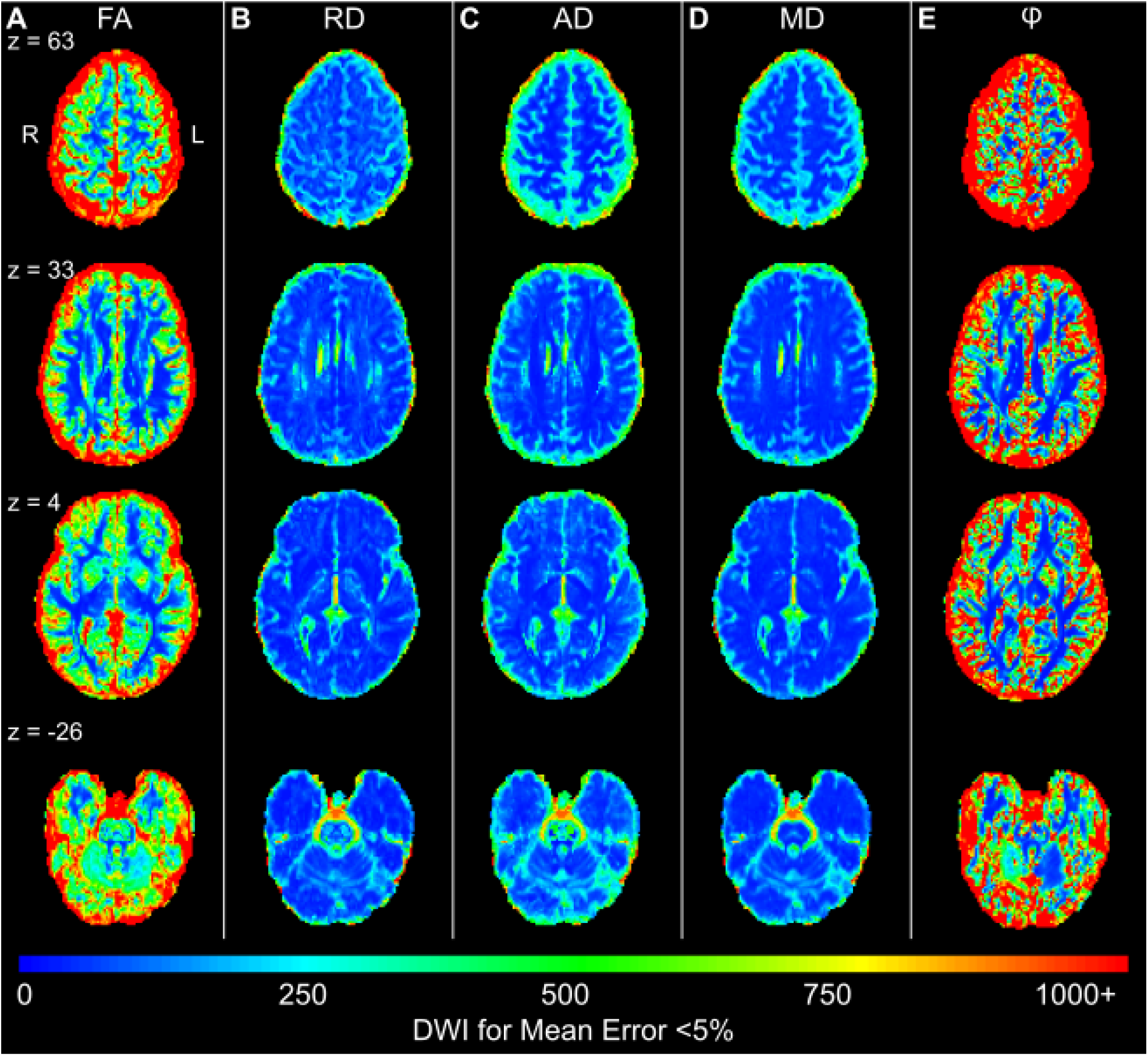
Whole-brain DTI Reliability Map (LLS) for Mean Error < 5% (Subject 2) (A) The color scale shows the number of DWI measurements needed to achieve a voxel-wise error less than 5% in FA. Error is calculated relative to the mean FA found using the entire sample. Results for (B) RD, (C) AD, (D) MD, and (E) angle φ are shown. Subjects 1 and 3 are shown in Figure S13

**Figure 7:**
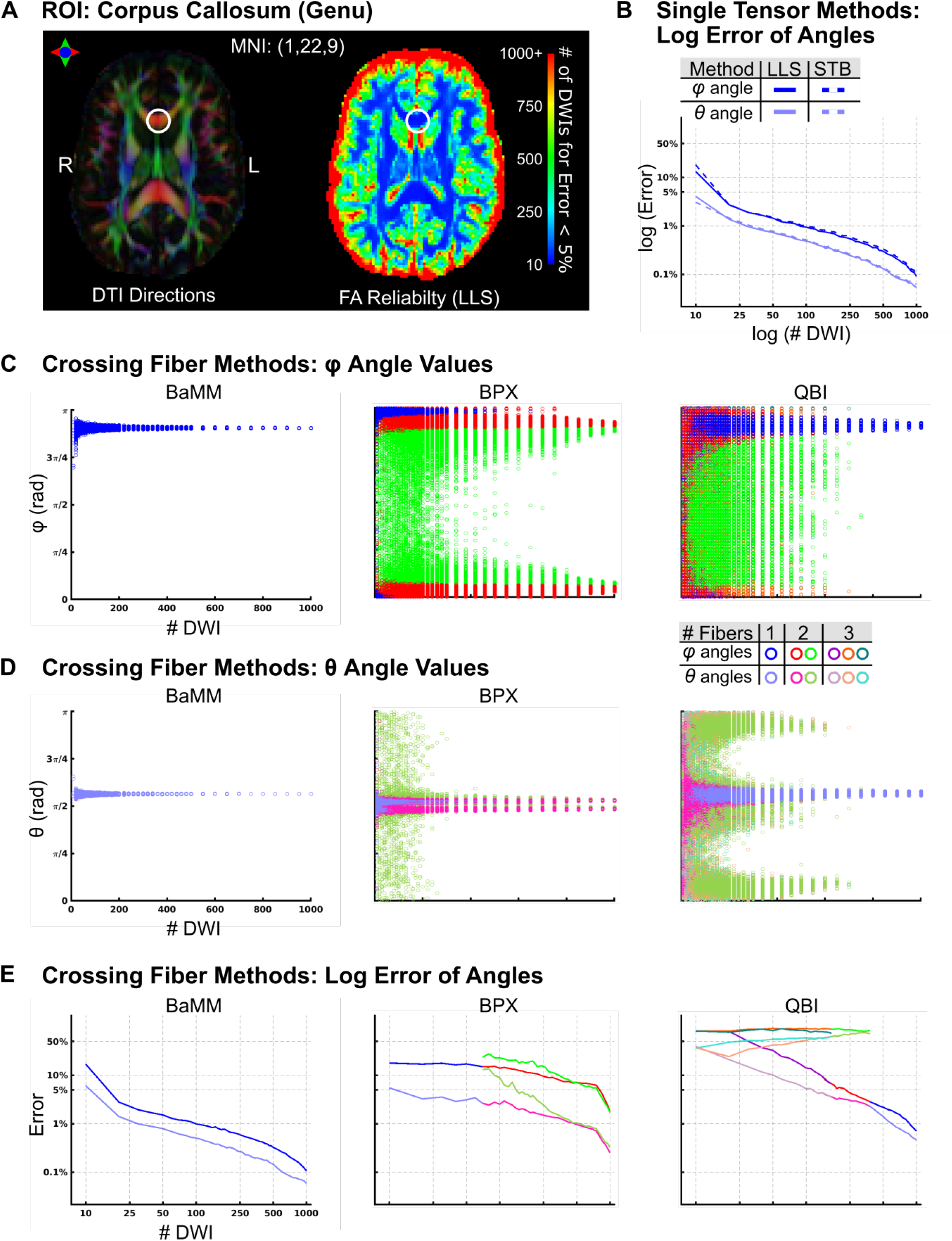
Reliability of Diffusion Measures in the Genu of the Corpus Callosum (Subject 2) (A) The locus of the analyzed voxel (MNI: 1, 22, 9) is marked with a circle. LLS FA reliability map as in Figure 6A. (B) Log Error of angle estimation by LLS and STB. (C) *φ* Angle estimations by Bayesian Multi-tensor Model-selection (BaMM), BedpostX (BPX), and Constant Solid Angle Q-Ball Imaging (QBI). (D) *θ* Angle estimations by BaMM, BPX, and QBI. (E) Log Error estimation by BaMM, BPX, and QBI. Error is calculated relative to the mean *φ* or *θ* found using the entire sample. Subject 1 and 3 in Figures S14 and S15, respectively

**Figure 8:**
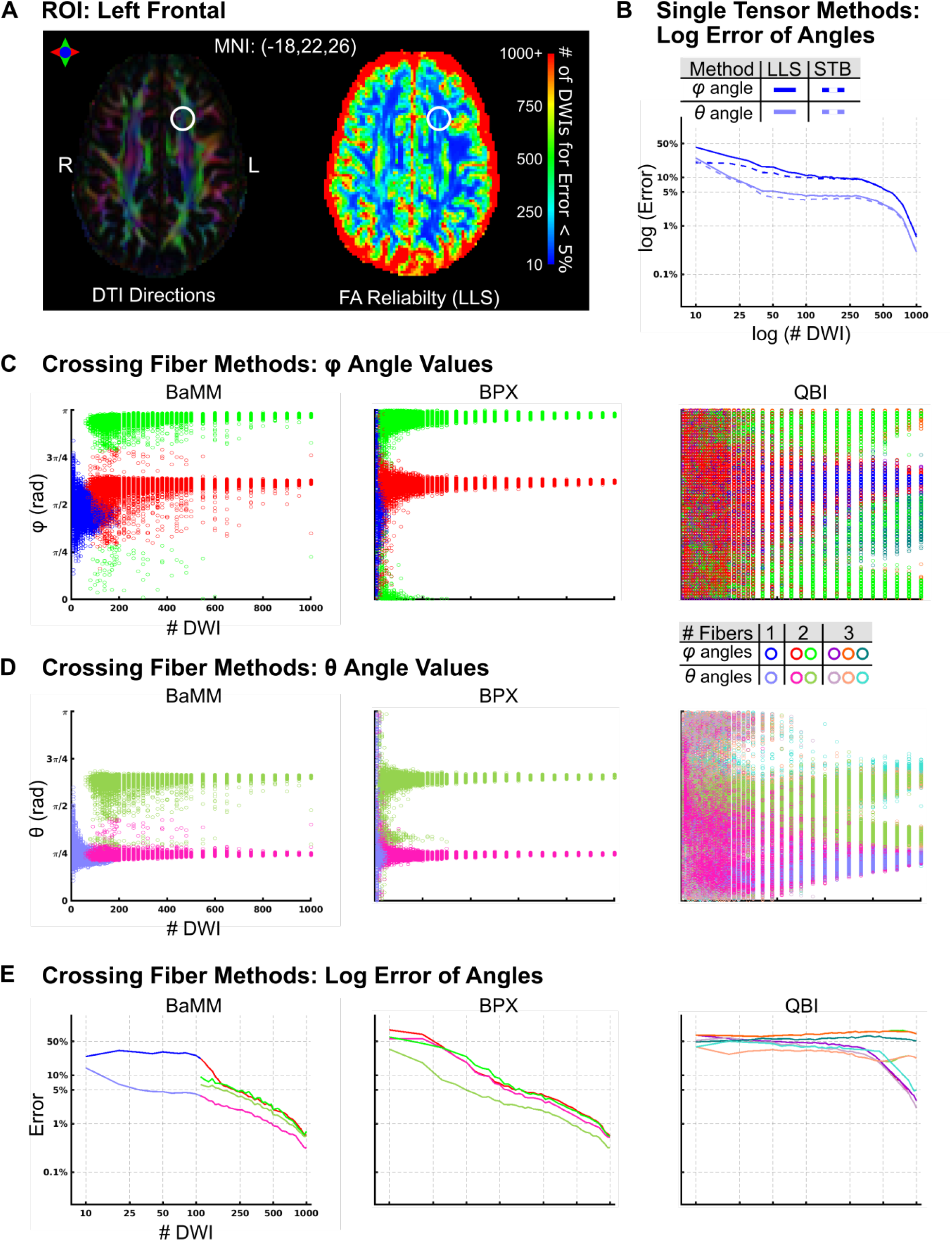
Reliability of Diffusion Measures in Left Frontal ROI (Subject 2) (A) The locus of the analyzed voxel (MNI: 18,22,26) is marked with a circle. LLS FA reliability map as in Figure 6A. (B) Log Error of angle estimation by LLS and STB. (C) *φ* Angle estimations by Bayesian Multi-tensor Model-selection (BaMM), BedpostX (BPX), and Constant Solid Angle Q-Ball Imaging (QBI). (D) *θ* Angle estimations by BaMM, BPX, and QBI. (E) Log Error estimation by BaMM, BPX, and QBI. Error is calculated relative to the mean φ or θ found using the entire sample. Subject 1 and 3 in Figures S16 and S17, respectively.

BaMM consistently and correctly estimated two fibers for > 30 DWI (red/pink and green/olive symbols Figure 4A-B, full parameter space in Figure S5). In contrast, BPX estimated two fibers at all but the smallest subsampling size (DWI = 10; Figure 4A-B, S6) when using default settings. When we increased BPX’s maximum allowable number of fibers to 3 (Figure S9), BPX frequently estimated three fibers for all DWI subsampling’s and the angle error increased in all three fibers when excess fibers were estimated.

Similar to the single tensor data, QBI incorrectly estimated three fiber directions at DWI < 150 (Figure 4A-B, S7). Even though two fiber directions were most commonly found at higher sampling density, some permutations still demonstrated three fiber directions at all subsampling sizes (62% at 100 DWI, 44% at 300 DWI, 33% at 500 DWI, 14% at 800 DWI).

### 3.3. Three Tensor Simulations: Accurate Estimates Achieved with Fewest DWIs using BaMM

The final simulated data we evaluated was of three crossing tensors, potentially observed in crowded areas of deep white matter, such as the corticospinal tract in the internal capsule. We explored the following forward-modeled parameter space similar to two-crossing tensors: fiber crossing angle (30°, 60°, 90°), relative weight of fiber compartments (33/33/33, 26/34/40, 13/34/53), SNR (30, 50, 100), and FA of tensors (0.6/0.6/0.6, 0.7/0.7/0.7, 0.8/0.8/0.8). Figure 5 shows the results of 90° crossing angle, 40/34/26 relative weight, SNR 50, and FA of 0.8/0.8/0.8. Results corresponding to the full parameter space are reported in the Supplemental Figures (Figures S10–12, BaMM, BPX, and QBI respectively). Results were consistent across most of the parameter space. All methods showed the lowest accuracy for the narrowest crossing fiber angle (30°).

BaMM consistently estimated three fiber compartments with sufficient data (DWI > 200, Figure 5A-B, S10). We increased BPX’s max possible fibers to 3 to match the simulated data, and then BPX estimated three fiber compartments at all subsampling sizes (Figure 5A-B, S11). As with prior modeling, BaMM and BPX correctly determined there were three fiber compartments and accurately estimated *ϕ* and *θ* for the three fibers with increasing sampling density. QBI most frequently estimated three fiber directions at all sampling densities, yet often estimated one or two fibers < 500 DWI (Figure 5A, S12).

For the three tensor simulations, BaMM and BPX log errors decreased almost linearly with an increase in the number of diffusion measurements (Figure 5C). QBI approached the expected relationship between log error and sample size once three fiber directions were consistently estimated (>500 DWI), yet still had higher error than BaMM and BPX at the largest subsampling sizes.

### 3.4. Whole Brain Reliability Mapping Reveals Very High Data Requirements in Gray Matter

To test reliability of diffusion metrics in human data (highly sampled, individual-specific), we used permutation subsampling of all available data to estimate whole-brain parameter variability (FA, RD, AD, MD, *ϕ, θ*), using the Linear Least Squares (LLS) method. LLS was used because none of the other methods were computationally tractable for whole brain analyses of this type, and whole brain reliability maps were desired to help identify anatomically defined regions of interest (ROI’s) (Figures 3–5). Figure 6 (and S13) shows the number of DWIs required to reach a mean error (RMSE) < 5% at each voxel. Error is now reported as the deviation from the mean when using the full sample (Eq. 4) rather than relative to the ground truth as in the prior simulations. AD, RD, and MD had less measurement error than FA and the angles *ϕ* and *θ* across most of the brain. In parts of the corpus callosum, only 20 DWIs were required for an FA RMSE < 5%. For most deep white matter voxels (e.g., corticospinal tracts, frontal crossing tracts), about 100 DWI samples were sufficient for an FA RMSE < 5%. In comparison, gray matter voxels required 300-500 measurements to reach an FA RMSE < 5%.

### 3.5. Corpus Callosum: Only BaMM Estimates Single Fiber < 600 DWIs

To examine individual-specific diffusion metric reliability with highly sampled data, across methods, several regions of interest (ROIs) were selected based on the whole-brain, voxel-wise LLS reliability maps (Figure 6) and prior anatomical knowledge. Figure 7 shows diffusion estimates in a voxel of the corpus callosum exemplifying highly anisotropic diffusion (Figure 7A; MNI: −1, 22, 9; Subject 2. This ROI in Subject 2 with BPX max 3 fibers in Figure S14. Subjects 1 and 3 in Figures S15–16 and S17, respectively): this ROI was chosen because it is strongly expected to contain only a single white matter fiber direction. Reliability curves for LLS and STB angles are shown in Figure 7B. As in the simulated single tensor data, these single tensor estimation methods showed low error rates (now reflecting reliability rather than accuracy), even for low DWI numbers.

Figure 7C-D shows the angle estimation results of the three more complex models: BaMM, BPX, and QBI. BaMM estimated only a single fiber in the corpus callosum, regardless of the number of DWIs in the subsample, with angles *φ* and *θ* closely matching the results observed with single tensor methods (see Figure 3, S3).

In contrast, BPX consistently estimated two fibers in the corpus callosum across all numbers of DWIs (unless otherwise specified, BPX results used default settings of max two fibers. BPX with max 3 fibers for this ROI in Figure S14). The principal fiber (red/pink) generally matched the orientation obtained with the other estimation methods (BaMM, QBI). At low sampling density, the angle of the second fiber estimated by BPX was broadly distributed over the interval 0 to *π* (green/olive). For DWI counts > 400, BPX continued to estimate two fibers, the average of which matched the orientation found by BaMM and QBI.

Similar to the simulated data, QBI estimated three fibers for subsamples with < 200 DWI, two fibers < 400 DWI, and a single fiber for > 400 DWI. For subsamples with < 200 DWI, QBI was most likely to estimate three fibers that were broadly distributed over the interval 0 to *π*, and also frequently estimated one or two fibers. Unlike BPX, QBI did consistently and accurately estimate one fiber for > 400 DWI. The existing anatomical priors about the corpus callosum would suggest a single primary diffusion direction, matching BaMM’s results at all subsampling sizes and QBI’s with ~ 1,000 DWIs.

### 3.6. Left Frontal White Matter: BPX with Two Fiber Default Setting Accurate with Fewest DWIs

We next selected a voxel in the left frontal lobe (MNI −18, 22, 26) where the superior longitudinal fasciculus and the uncinate fasciculus cross (Figure 8A). This voxel was chosen to be > 10 mm from any gray matter voxel in all three subjects (subjects 1 and 3 in Figures S18 and S19, respectively). Reliability curves for LLS and STB angle metrics (*ϕ, θ*) are shown in Figure 8B as controls. The single tensor models are inadequate to describe the full microstructural complexity, and increased error can be observed in Figure 8B vs Figure 7B.

Figure 8C-D contrasts the angle measurement reliability of the crossing fibers models (BaMM, BPX, and QBI). For very low numbers of DWI per subsample (< 50), BaMM identified the principal diffusion direction, whereas BPX returned approximately uniform density of diffusion directions at all angles (i.e., little to no angular information).

BaMM consistently estimated two diffusion directions with > 100 DWI. BPX consistently estimated two directions with > 20 DWI. Angular measurement error was generally less with BPX than BaMM, but comparable for > 250 DWI.

QBI most frequently estimated three fiber directions at all subsampling sizes, but also frequently estimated one or two fibers. The QBI estimation of two or three fibers was broadly distributed over 0 to *π* for < 500 DWI, and the error only improved marginally with increasing DWIs.

#### 3.6.1. Right Corticospinal Tract: Poor Reliability, BaMM Estimates Relatively Superior

The third ROI we analyzed in depth was in the right corticospinal tract as it progressed through/near the internal capsule, a brain region with potentially three crossing fibers (MNI 22, −19, 11; Figure 9A). Based on anatomical priors, model sensitivity, registration to MNI coordinates, and accuracy of ROI location across subjects, we could expect a single fiber direction reflecting the CST, two fiber directions for the CST and internal capsule, or three directions for a fanning behavior of either the CST or internal capsule fibers. BPX settings were set to a maximum of three fibers accordingly. Results for subjects 1 and 3 are in Figures S20 and S21, respectively. Control reliability curves for LLS and STB angle metrics (*ϕ, θ*) are shown in Figure 9B and demonstrate the expected increase in reliability as a function of DWI number.

**Figure 9:**
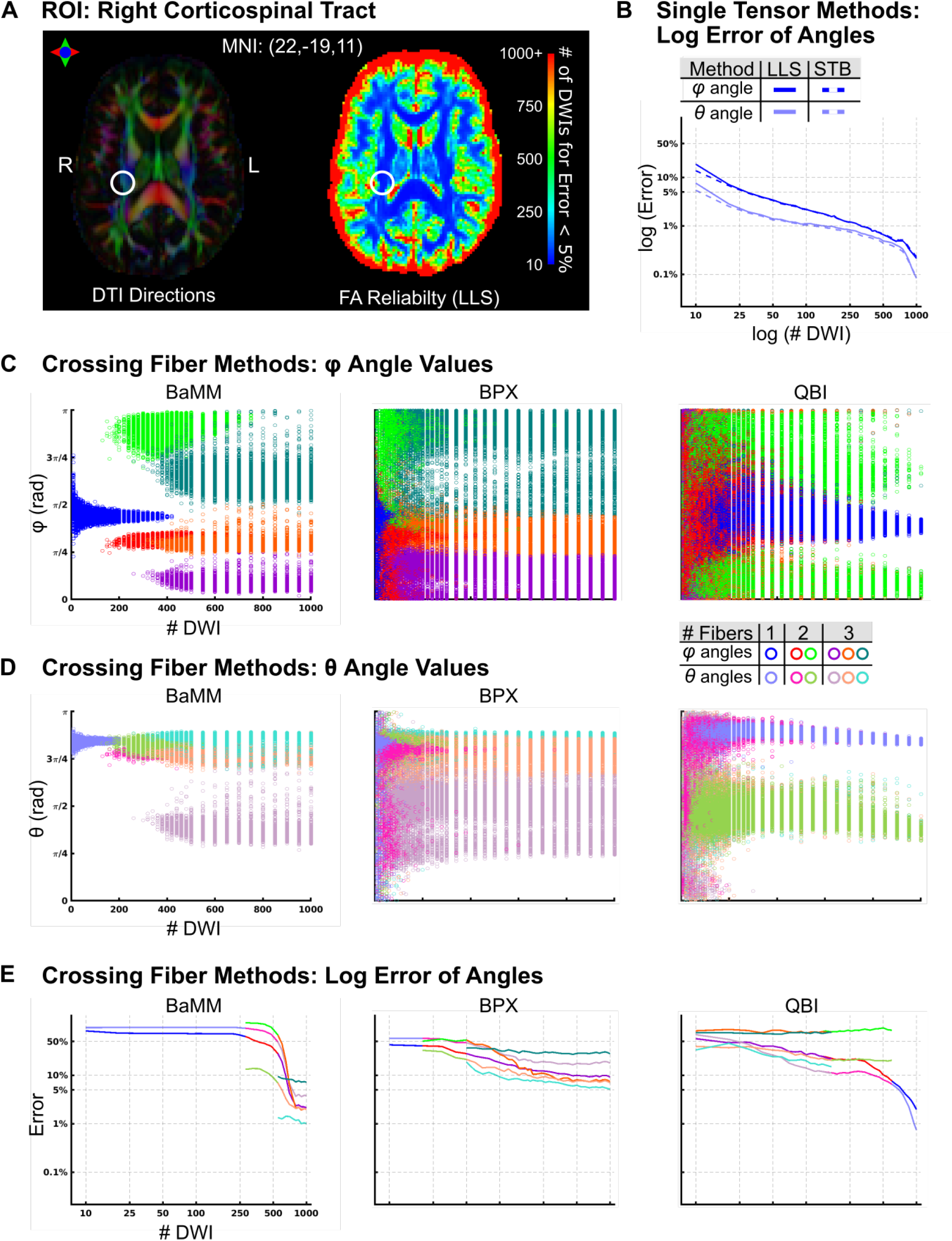
Reliability of Diffusion Measures in Right Corticospinal Tract (Subject 2) (A) The locus of the analyzed voxel (MNI: −22, −19,11) is marked with a circle. LLS FA reliability map as in Figure 6A. (B) Log Error of angle estimation by LLS and STB. (C) *φ* Angle estimations by Bayesian Multi-tensor Model-selection (BaMM), BedpostX (BPX), and Constant Solid Angle Q-Ball Imaging (QBI). (D) *θ* Angle estimations by BaMM, BPX, and QBI. (E) Log Error estimation by BaMM, BPX, and QBI. Error is calculated relative to the mean *φ* or *θ* found using the entire sample. Subject 1 and 3 in Figures S18 and S19, respectively.

Figure 9C-D contrasts the angle measurement reliability of the crossing fibers models (BaMM, BPX, and QBI). For samples with < 260 DWIs, BaMM estimated a single fiber; between 260 and 500 DWI BaMM most frequently estimated two fibers; and for DWIs > 500, BaMM estimated three fibers. However, the distribution of angle estimates (*φ, θ*) was broader than seen in previous ROIs. With > 50 DWIs, BPX estimated three fibers with almost uniform distribution of *ϕ* and *θ* from 0 to *π*. In contrast, QBI followed an estimation pattern previously observed in the corpus callosum and simulated single tensor data (Figures 3 and 7). For < 200 DWI QBI estimated three fibers, then two fibers for < 550 DWI, and for > 550 DWI a single primary fiber was estimated.

Error estimation (relative to the mean) improved with increasing number of DWI for all methods. BaMM’s error estimation stabilized once the model selection consistently chose three fibers (Figure 9E, >500 DWI). However, estimated error in this ROI was higher than in previously tested ROI. BPX error estimation also improved with increasing sample size but remained above 5% for all subsampling sizes. Similarly, QBI estimated error decreased for the primary fiber direction with increasing sample size. However, since the three models diverged in their estimation of number of fibers and the orientation of those fibers, we can only speak to the reliability of the models and not their accuracy.

## 4. DISCUSSION

Identifying and understanding inter-individual differences in brain organization is critically important for neuroscience, neurology, neurosurgery, and psychiatry [43]. While almost all typically developing individuals share the same major white matter bundles [2], variations in size, position, and/or orientation of white matter fibers could have significant effects on surgical plans, and recovery from brain injury. In addition, individual-specific precision diffusion imaging is necessary to effectively bridge functional and structural connectivity. To fully capitalize on the microstructural information revealed by DTI and other methods, we must improve our ability to generate precise, individual-specific diffusion imaging maps.

### 4.1. BaMM: A Novel Estimation Method for Diffusion Imaging Robust to Overfitting

Accuracy is inherently limited in single-tensor models because modeling complex white matter microstructure as a single tensor may obscure valuable information, i.e., crossing fibers. While existing techniques can model multiple fibers, they are limited because of the model’s shape assumptions. Therefore, we developed a novel Bayesian Multi-tensor Model-selection (BaMM) method that can precisely and accurately fit single- and crossing-tensor DWI data (simulated; highly sampled, individual-specific). The novel BaMM parameter estimation algorithm was based on previous Bayesian, overfitting robust modelselection work [44]. In particular, we designed BaMM to compare multiple models and select the model best suited for the available data. Here, we demonstrated that BaMM can accurately estimate one, two, or three tensors, and that its precision improves with increasing DWI number. In the current implementation, BaMM uses the same assumed model as BPX (ball-and-sticks; [10]) but can accommodate a large input data set by scaling the parameter estimation penalties to the dataset size. We tested and validated BaMM over a wide range of diffusion measurements (10 to 1000) to rule out bias for a specific number of DWIs. The current work was completed using a ball-and-sticks partial volumes model, but BaMM can also accommodate full multi-tensor models, multi-fiber kurtosis models, or other models yet to be developed [13, 45]. The BaMM framework is adaptable to any set of mathematical assumptions about white matter structure and can serve to directly compare different diffusion models.

### 4.2. BPX: Accurate and Reliable if Assumptions Are Met

BPX inspired the creation of BaMM but differs in its parameter estimation approach. BPX was initially optimized for about 30 DWI, and then updated for 60 DWI [11, 46]. The datasets analyzed here contained much higher quantities of high-quality data (800+ low-motion DWIs for each individual). With an excess of DWIs, we observed that BPX consistently estimates the maximum allowed number of fibers. BPX’s default setting is a maximum of two fibers, and with these settings BPX estimated two fibers for simulated single tensor data (Figure 3) and for the corpus callosum (Figure 7). When set to allow for a maximum of three tensors, BPX estimated three fibers in simulated single tensor data and in the corpus callosum ROI (Figure S4 and S14). When provided with a large number of DWIs (> 500), BPX’s splitting of a single tensor into two that are almost superimposed is inaccurate but not detrimental to subsequent tractography. By contrast, a potentially inappropriately oriented second or third fiber, could substantially deviate probabilistic tracking (Fig 9C-D). Nevertheless, we note that when the data match BPX’s assumptions, it is accurate and reliable from 10-1,000 DWIs. Determining the appropriate priors for all voxels in the brain, however, is a significant challenge.

### 4.3. QBI: Accurate Only with Very Large Amounts of Diffusion Data

Constant solid angle Q-Ball Imaging (QBI) was designed to eliminate diffusion tensor shape assumptions, and it estimates water diffusion using spherical harmonics [12, 35]. QBI can estimate one, two or three tensors with 1,000 DWIs, but problematically, the reliability of these estimates always remained low (Fig 3–5, 7–9). With < 800 DWIs QBI tends to model additional fiber directions, possibly capturing noise in the data. Since the published literature recommends a specific acquisition scheme sequence for QBI (i.e., high angular resolution; at least 500 DWI) [12], we collected supplementary DWI data with 960 unique B-vectors and 50 B0 (1020 DWI). Supplemental analyses (Figure S16) revealed that none of the tested methods, QBI included, improved with high angular resolution data. Instead, it appears that repeated acquisitions of ABCD’s 103 DWI protocol were less prone to overfitting [18]. Combining DWI samples over multiple sessions introduces jitter in angular sampling owing to variability of head position, and effectively provides the needed angular resolution.

### 4.4. Reliability of Diffusion/Anisotropy and Fiber Orientation Increases with Increasing DWI Number

In both highly sampled human and simulated data, estimate variability decreased with increasing sample size. Standard error is defined as the standard deviation divided by the square root of the sample size, therefore assuming a normal distribution measurement error should be inversely related to the square root of the sample size. Our results failed to follow this pattern under two conditions: when there was insufficient data to constrain the model (e.g., < 20 DWI for LLS and STB), or when the model misrepresented the underlying diffusion process (i.e., using single tensor methods for multiple fibers, or assuming multiple fibers for a single tensor). Overall, deep white matter voxels demonstrated lower measurement error than the rest of the brain, and larger data amounts were needed for voxels with lower FA (Figure 6). FA measurement error was < 5% with 70-150 DWIs in deep white matter, while cortical voxels required 300-500 DWIs to comparably reduce error. Angles *ϕ* and *θ* showed the highest measurement error of all the diffusion metrics. Uncertainty in the angle of the tensor is related to uncertainty in anisotropy, explaining why angle error estimation is higher in gray matter [16].

### 4.5. Precision Diffusion Imaging is Achievable with Realistic Data Acquisition Times

Our work demonstrates the feasibility of implementing individual-specific, precision diffusion imaging. Only 15 to 30 DWIs commonly are acquired in clinical settings, requiring very short scan times of < 1 minute. As MRI hardware and processing software improved, researchers have started to acquire larger diffusion data sets (100 - 300 DWIs per subject) while maintaining reasonable imaging times (6 - 20 min). Our study demonstrated that one can reliably estimate the shape and orientation of a single diffusion tensor in deep white matter with about 100 diffusion measurements. Thus, not only research DTI scans [17, 18, 47] but also clinical ones should collect a greater number of DWIs (at least ~100).

For crossing-fiber diffusion models, at least 300 DWIs are generally required in deep white matter, assuming high data quality. To advance from 100 to 300 DWI requires an increase in total scan time from about 6 minutes to about 20 minutes. Acquiring 1,000 DWIs takes a little over an hour. An hour-long diffusion scan can may be warranted for individual-specific precision mapping for research or in neurosurgical planning [48]. Diffusion data acquisition is typically better tolerated than task or resting state functional MRI (fMRI) because the patient can sleep or watch a movie during the scan. Therefore, a small additional investment in scanning time could have significant positive effects on diagnostics and treatment of individual neurological and neurosurgical patients. In addition, acquiring greater amounts of high-quality DTI data would expand the available processing schemes beyond the models described here to methodologies which are even data hungrier (e.g., DSI, DBSI) [14, 47].

### 4.6. Precision Diffusion Imaging to Enhance Structural Connectivity Maps (End-to-End Tracking) in Cortex

Researchers have been exploring the feasibility and validity of MRI based structural connectivity analyses since before the export of these techniques to fMRI data. [41, 49–55]. Many studies that attempt to build structural connectivity maps initiate the fiber tracking at the border of gray and white matter. Since FA and angle orientations (*ϕ* and *θ*; Figure 6, S13) are less reliable closer to gray matter, more errors are introduced at initiation of the tracking. Though many other challenges to structural connectivity maps must still be addressed [56, 57], structural connectivity and other advanced modeling techniques would likely also greatly benefit from larger numbers of DWIs per individual.

Individual-specific precision DTI will facilitate and enrich clinical and investigative neuroimaging. Higher-fidelity DTI holds great promise for surgical planning and post-surgical evaluation (e.g., epilepsy disconnection procedures [58]), and for any studies seeking to evaluate longitudinal changes within-subject (ABCD; [59]). As with precision functional mapping (PFM), we cannot fully predict the fine-grained details and individual-specific variants that precision DTI will reveal, but precise individual-specific structural connectomes will very likely increase our understanding of brain architecture in general, and of individual variability in health and disease, in particular.

## Acknowledgements

This work was supported by NIH grants NS088590 (N.U.F.D.), MH96773 (N.U.F.D.), MH122066 (N.U.F.D.), MH124567 (N.U.F.D.), MH121276 (N.U.F.D.), HD087011 (J.S.S.), NS115672 (A.Z.), NS110332 (D.J.N.), MH1000872 (T.O.L.), MH112473 (T.O.L.), NS090978 (B.P.K.), MH121518 (S.M.), MH104592 (D.J.G.), HD094381 (Y.W.), AG053548 (Y.W.), NS098577 (to the Neuroimaging Informatics and Analysis Center); the US Department of Veterans Affairs Clinical Sciences Research and Development Service grant 1IK2CX001680 (E.M.G.); Kiwanis Neuroscience Research Foundation (N.U.F.D.); the Jacobs Foundation grant 2016121703 (N.U.F.D.); the Child Neurology Foundation (N.U.F.D.); the McDonnell Center for Systems Neuroscience (D.J.N., T.O.L., A.Z., B.L.S., and N.U.F.D.); the McDonnell Foundation (S.E.P.), the Mallinckrodt Institute of Radiology grant 14-011 (N.U.F.D.); the Hope Center for Neurological Disorders (N.U.F.D., B.L.S., and S.E.P.); the Intellectual and Developmental Disabilities Research Center at Washington University (J.S.S); the Bright-Focus Foundation grant A2017330S (Y.W.); the March of Dimes (Y.W.).

## Supplemental Materials

**Figure 1:**
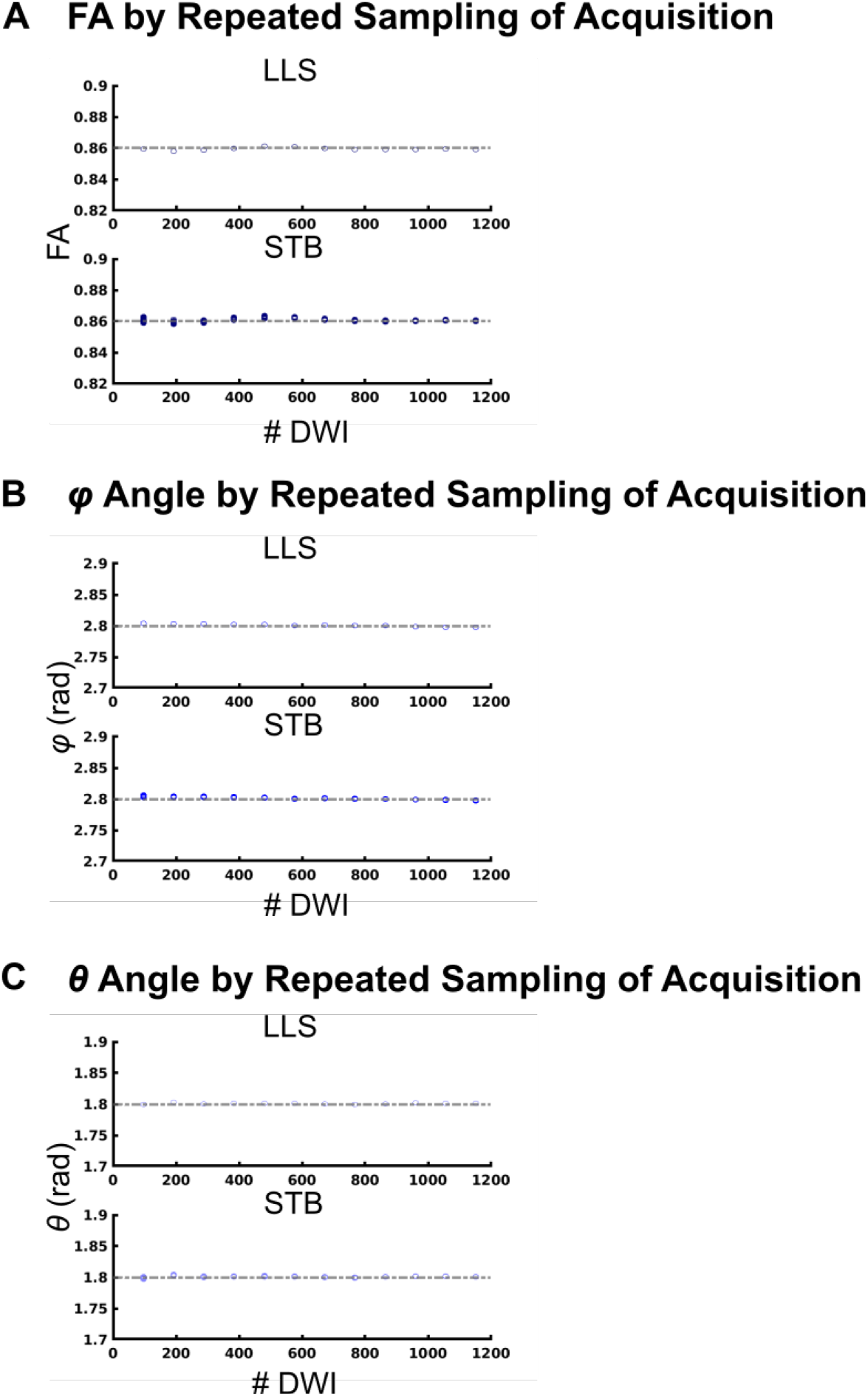
Validation of Subsampling Repeated sampling selecting an entire acquisition of 103 DWI (so select 1-12 acquisitions). (A) FA by LLS and STB with repeated sampling of 1-12acquisitions. (B) Angle *φ*(C) Angle *θ*

**Figure 2:**
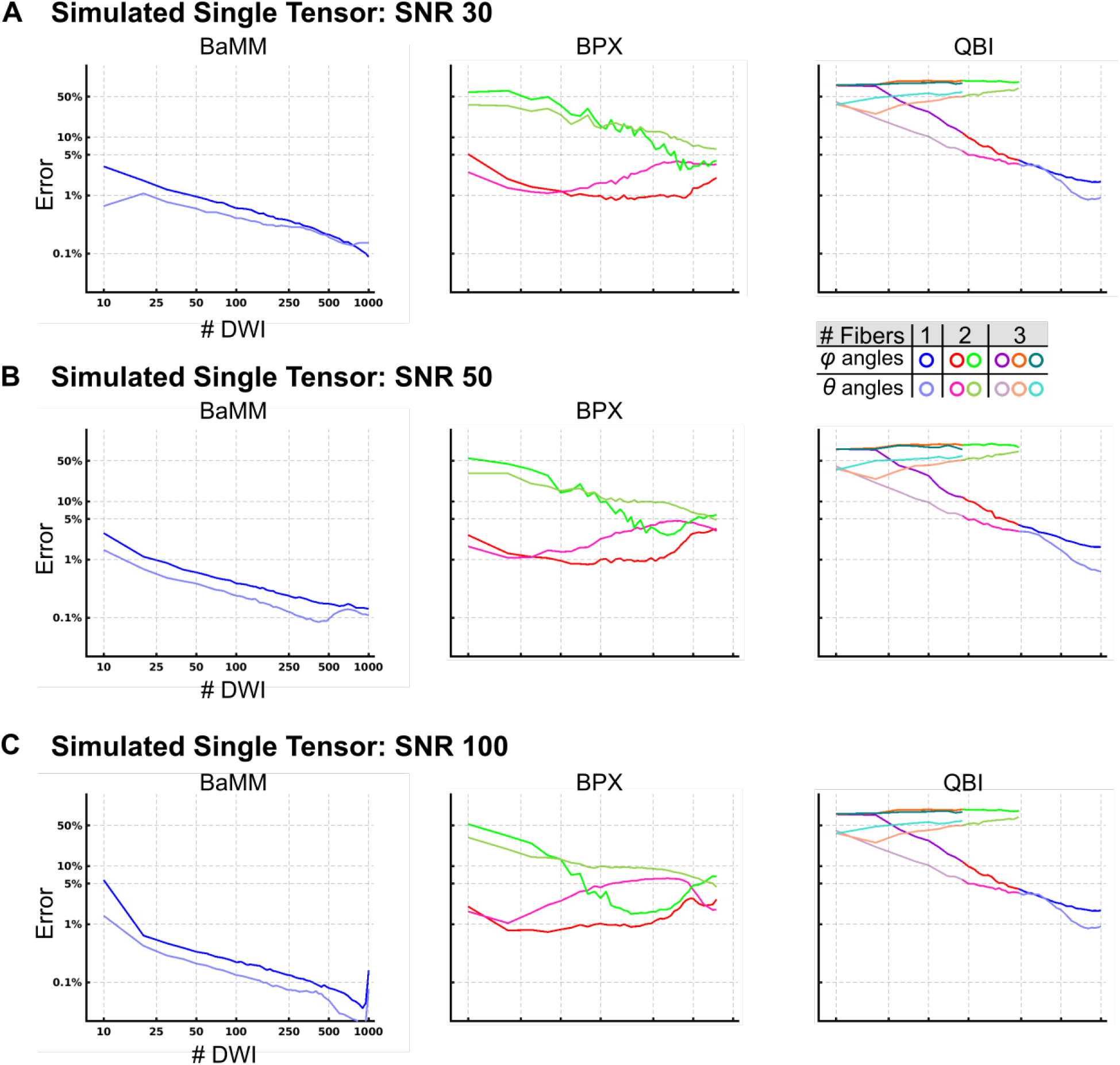
Accuracy of DTI Measures by BaMM, BPX, QBI, Simulated Single Tensor (A)Error estimation by Bayesian Multi-tensor Model-selection (BaMM), BedpostX (BPX), and constant solid angle Q-Ball Imaging (QBI) at SNR30. Mean error at each subsampling size was calculated, then plotted on a log scale. Plots are colored by the most frequent number of fibers estimated: subsamples with a single fiber direction are plotted in blue/sky blue, two fibers plotted red/pink and green/olive, 3 fibers are plotted in purple/lilac, orange/salmon, teal/cyan. (B) SNR50. (C) SNR 100.

**Figure 3:**
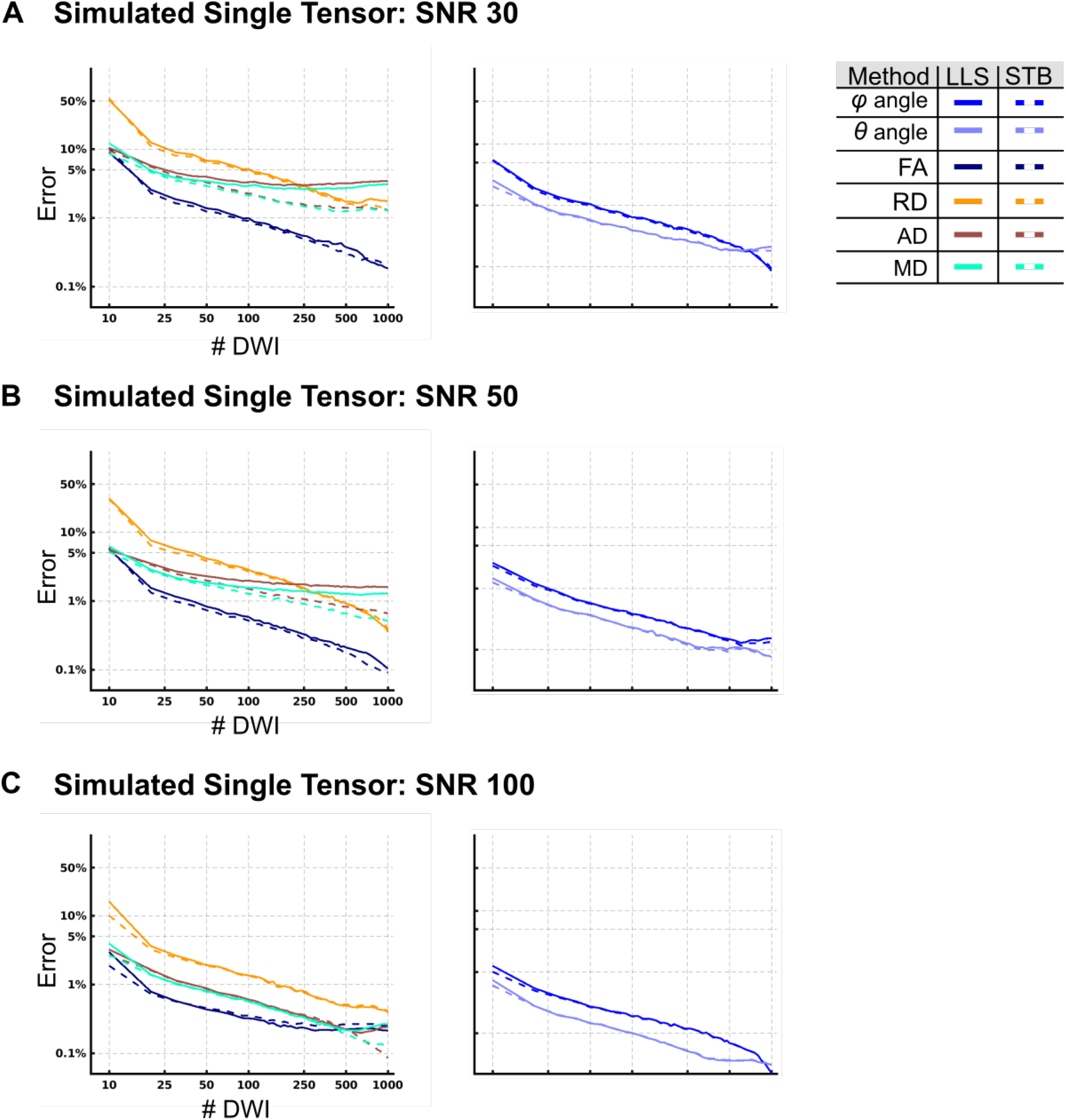
Accuracy of DTI Measures by LLS and STB, Simulated Single Tensor (A) Error estimation by Linear Least Squares (LLS) and Single TensorBayesian (STB) at SNR30. Mean error at each subsampling size was calculated, then plotted on a log scale. LLS and STB plotted using straight and dotted lines, respectively. FA is in navy, RD in yellow, AD in brown, and MD in bright green. *φ*is plotted in blue, and *θ* sky blue. (B) SNR 50. (C) SNR 100

**Figure 4:**
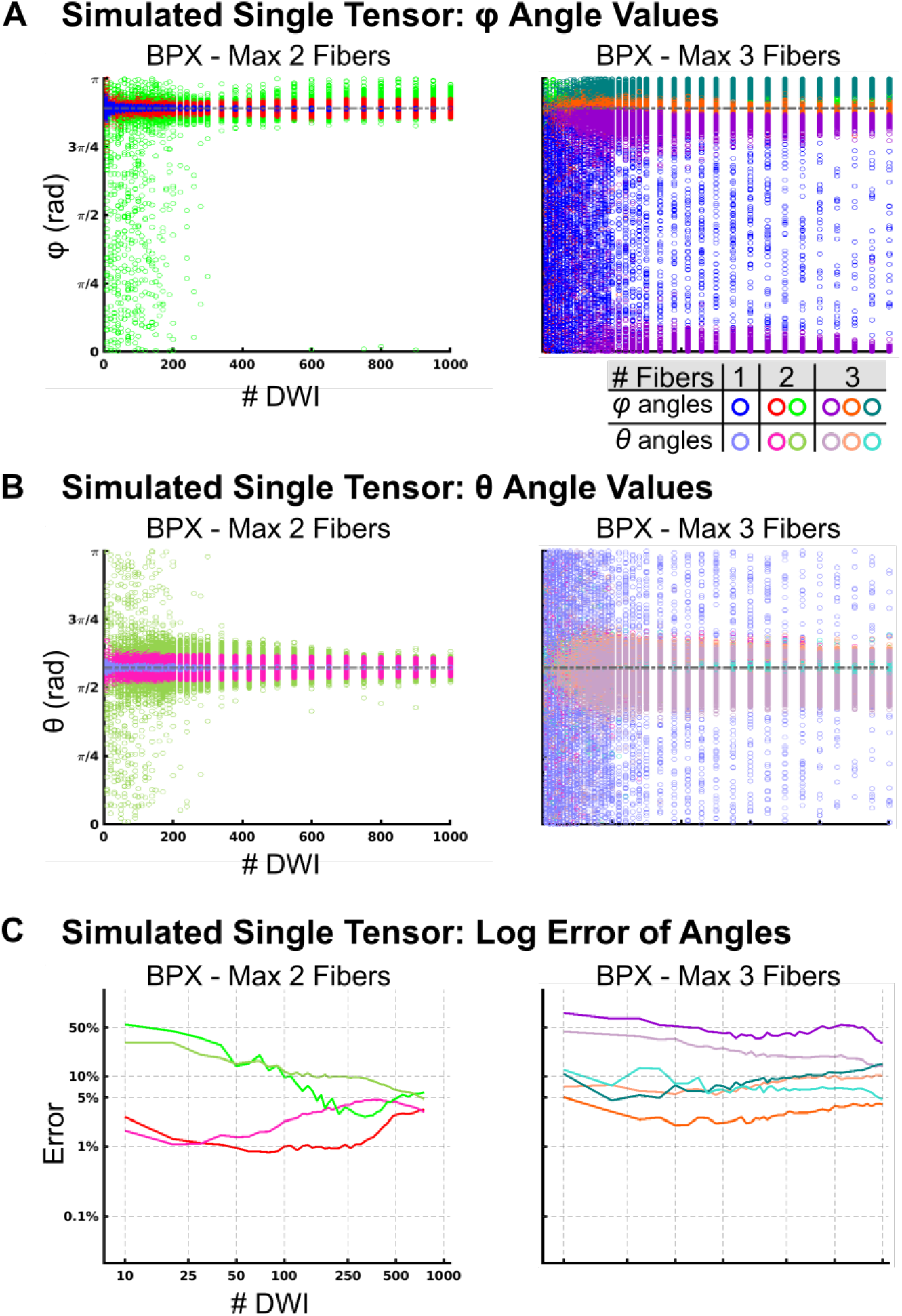
Accuracyof DTI Measures by BPX, Simulated Single Tensor with Max 3 Fibers (A) *φ* angle estimations by BedpostX (BPX) with maximum2 or 3 fibers. Open circles represent the results obtained by repeated permutation sampling. Same color legend for all data panels. Permutations that resulted in a single fiber direction are plotted in blue (φ). Permutations that resulted in two fibers are plotted in red (*φ*) and green (*φ*). Permutations that resulted in three fibers are plotted in purple (*φ*), orange (*φ*), teal (*φ*). (B) *θ* angle estimations. Permutations are plotted in sky blue (*θ*) for one fiber, pink(*θ*) and olive (*θ*) for two fibers, and lilac (*θ*), salmon (*θ*), cyan (*θ*) for three fibers. (C) Error estimation for BPX with max 2 or 3 fibers. Mean error at each subsampling size was calculated, then plotted on a log scale. The same colors as in (A/B) are used and indicate the most frequent number of fibers estimated at each subsampling size

**Figure 5:**
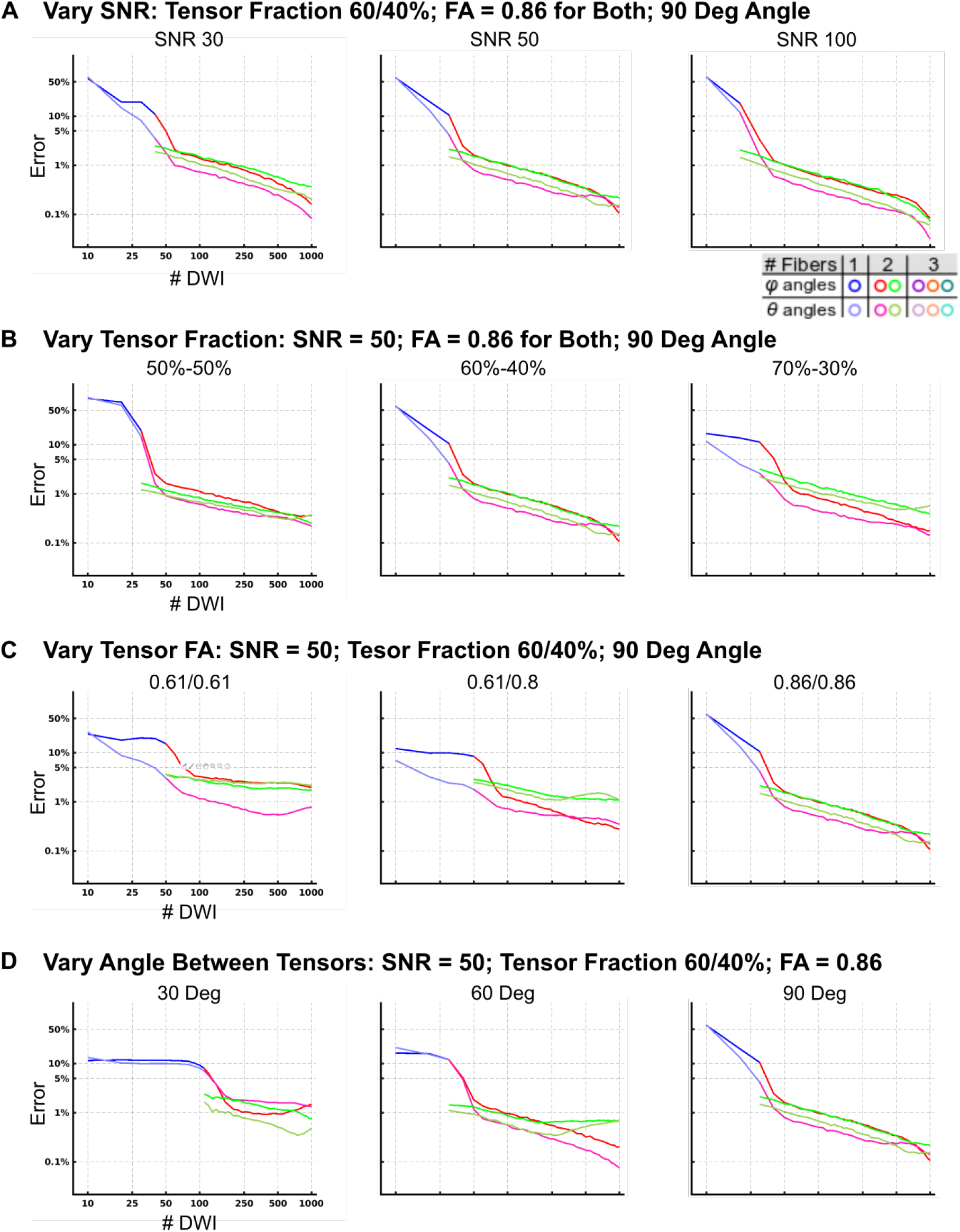
Accuracy of BaMM in variety of two tensor simulated data (A)Error estimation by BaMM for two crossing tensors at varying SNR. Mean error at each subsampling size was calculated, then plotted as the log of error. Plots are colored by the most frequent number of fibers estimated: subsamples with a single fiber direction are plotted in blue/sky blue, two fibers plotted red/pink and green/olive, 3 fibers are plotted in purple/lilac, orange/salmon, teal/cyan. (B) Vary tensor Fraction. (C) Vary tensor FA. (D) Vary angle between tensors.

**Figure 6:**
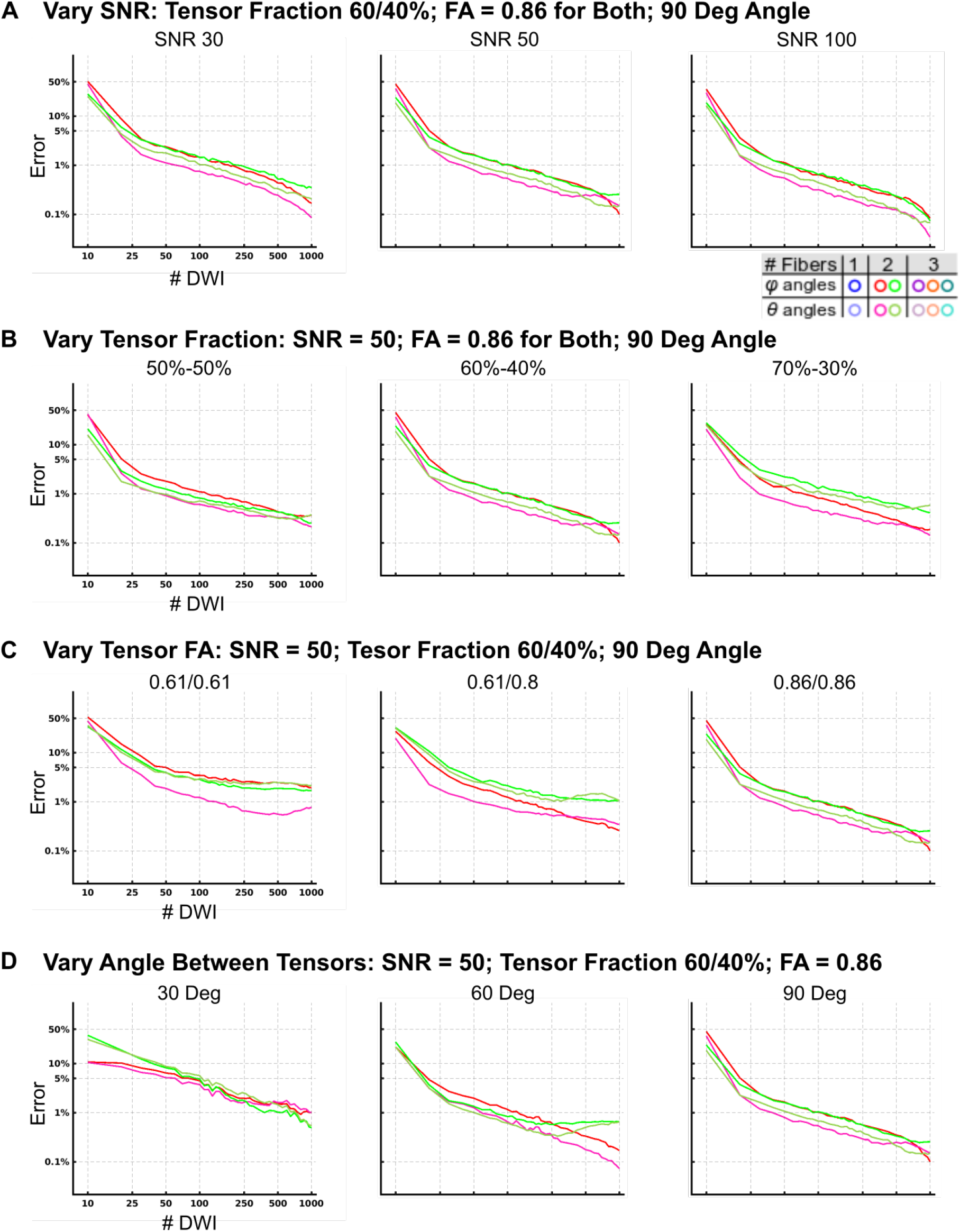
Accuracy of BPX in variety of two tensor simulated data (A) Error estimation by BPX for two crossing tensors at varying SNR. Mean error at each subsampling size was calculated, then plotted as the log of error. Plots are colored by the most frequent number of fibers estimated: subsamples with a single fiber direction are plotted in blue/sky blue, two fibers plotted red/pink and green/olive, 3 fibers are plotted in purple/lilac, orange/salmon, teal/cyan. (B) Vary tensor Fraction. (C) Vary tensor FA. (D) Vary angle between tensors.

**Figure 7:**
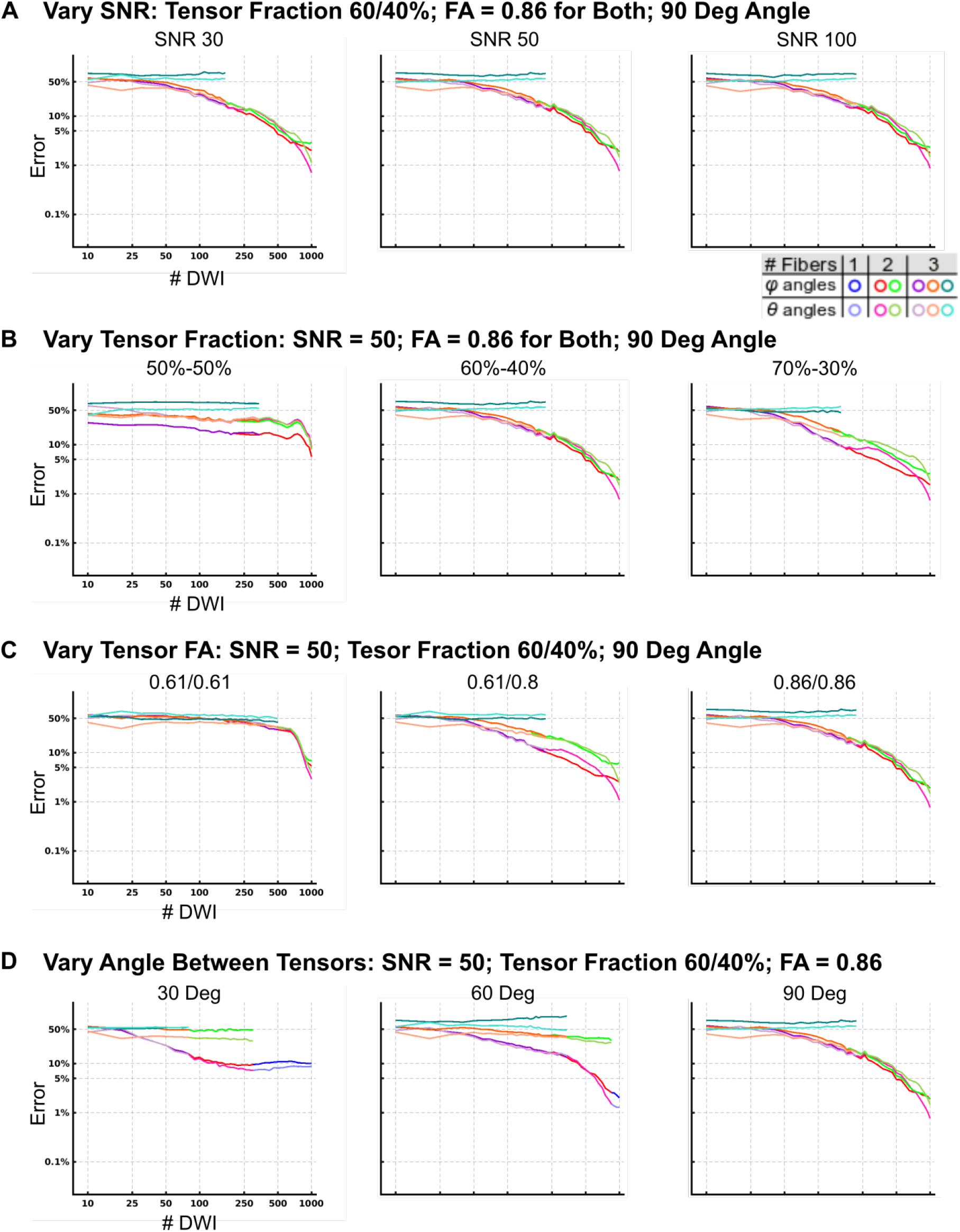
Accuracy of QBI in variety of two tensor simulated data (A) Error estimation by QBI for two crossing tensors at varying SNR. Mean error at each subsampling size was calculated, then plotted as the log of error. Plots are colored by the most frequent number of fibers estimated: subsamples with a single fiber direction are plotted in blue/sky blue, two fibers plotted red/pink and green/olive, 3 fibers are plotted in purple/lilac, orange/salmon, teal/cyan. (B) Vary tensor Fraction. (C) Vary tensor FA. (D) Vary angle between tensors.

**Figure 8:**
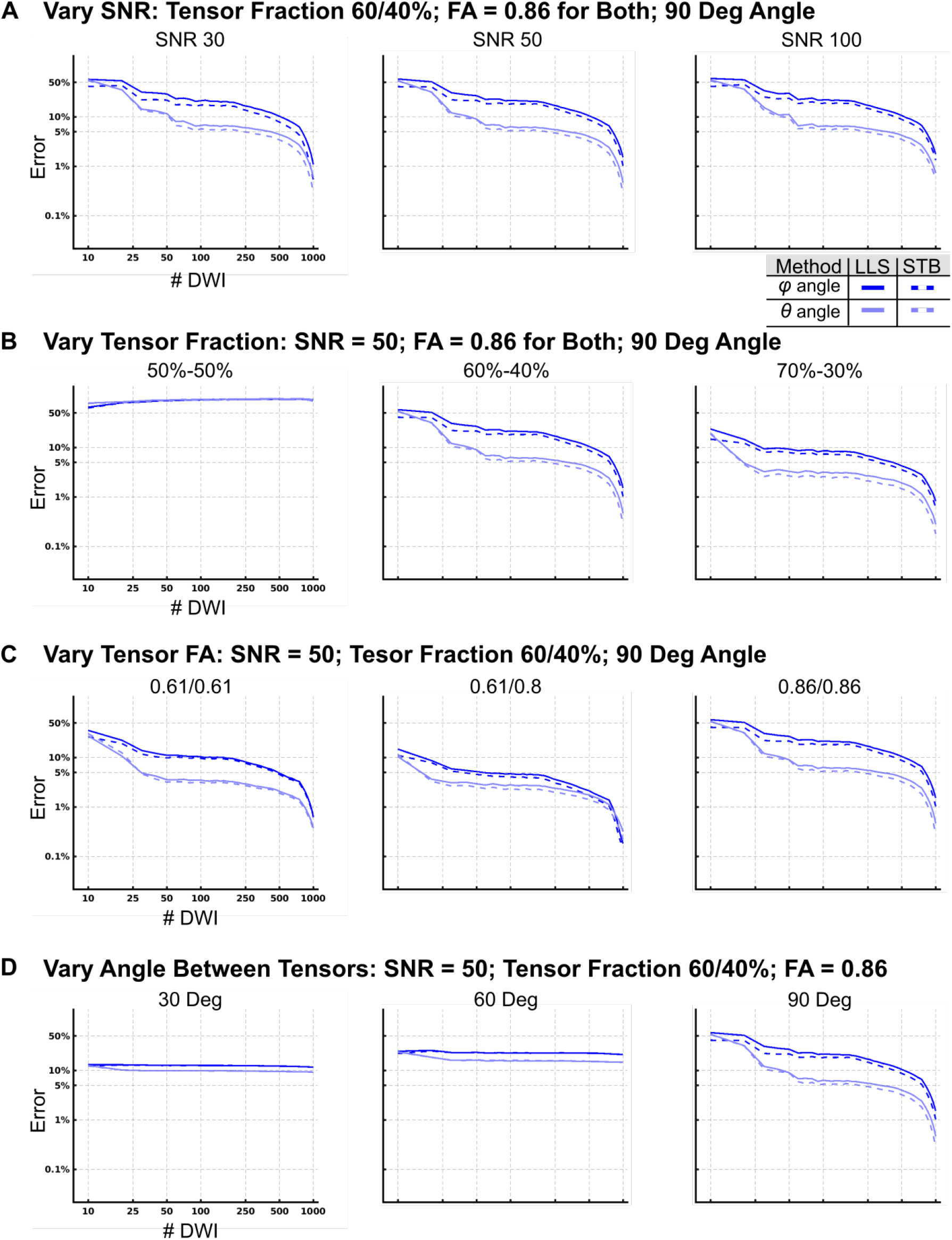
Accuracy of LLS and STB in variety of two tensor simulated data (A) Error estimation by LLS and STB for two crossing tensors at varying SNR. Mean error at each subsampling size was calculated, then plotted as the log of error. Plots are colored by the most frequent number of fibers estimated: subsamples with a single fiber direction are plotted in blue/sky blue, two fibers plotted red/pink and green/olive, 3 fibers are plotted in purple/lilac, orange/salmon, teal/cyan. (B) Vary tensor Fraction. (C) Vary tensor FA. (D) Vary angle between tensors.

**Figure 9:**
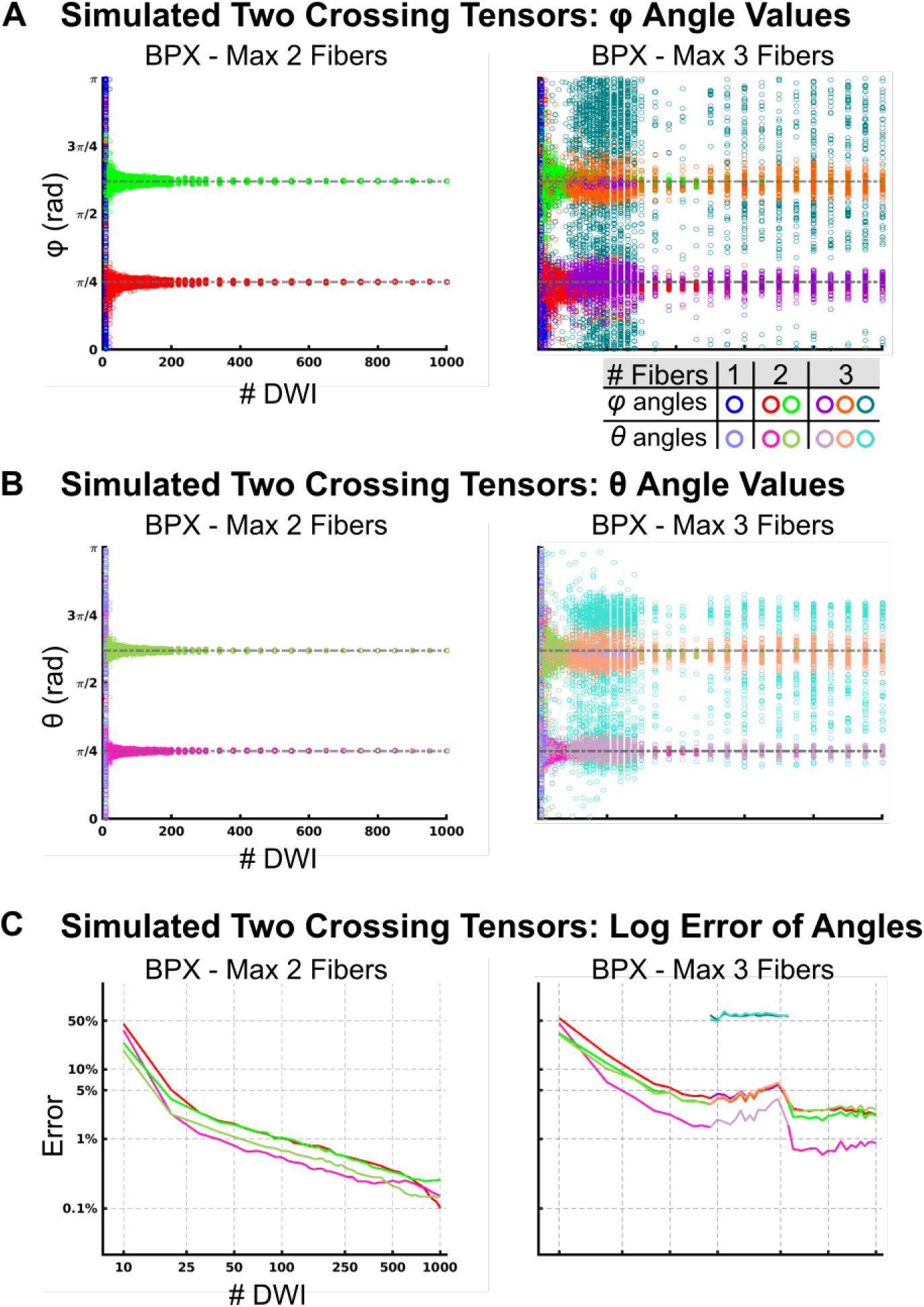
Accuracy of BPX in two tensor simulated data with max 3 fibers The tensors were oriented such that they were perpendicular to each other. The first tensor had larger weighting equal to 60% of the signal. Rician noise was added for an SNR = 50. (A) *φ* angle estimations by BedpostX (BPX), with max 2 or three fibers. Open circles represent the results obtained by repeated permutation sampling. Same color legend for all data panels. Permutations that resulted in a single fiber direction are plotted in blue (*φ*). Permutations that resulted in two fibers are plotted in red (*φ*) and green (*φ*). Permutations that resulted in three fibers are plotted in purple(*φ*), orange (*φ*), teal (*φ*). (B) *θ* angle estimations. Permutations are plotted in sky blue (*θ*) for one fiber, pink (*θ*) and olive (*θ*) for two fibers, and lilac(*θ*), salmon (*θ*), cyan (*θ*) for three fibers. (C) Error estimation for BPX with max two or three fibers. Mean error at each subsampling size was calculated, then plotted on a log scale. The same colors as in (A/B) are used and indicate the most frequent number of fibers estimated at each subsampling size.

**Figure 10:**
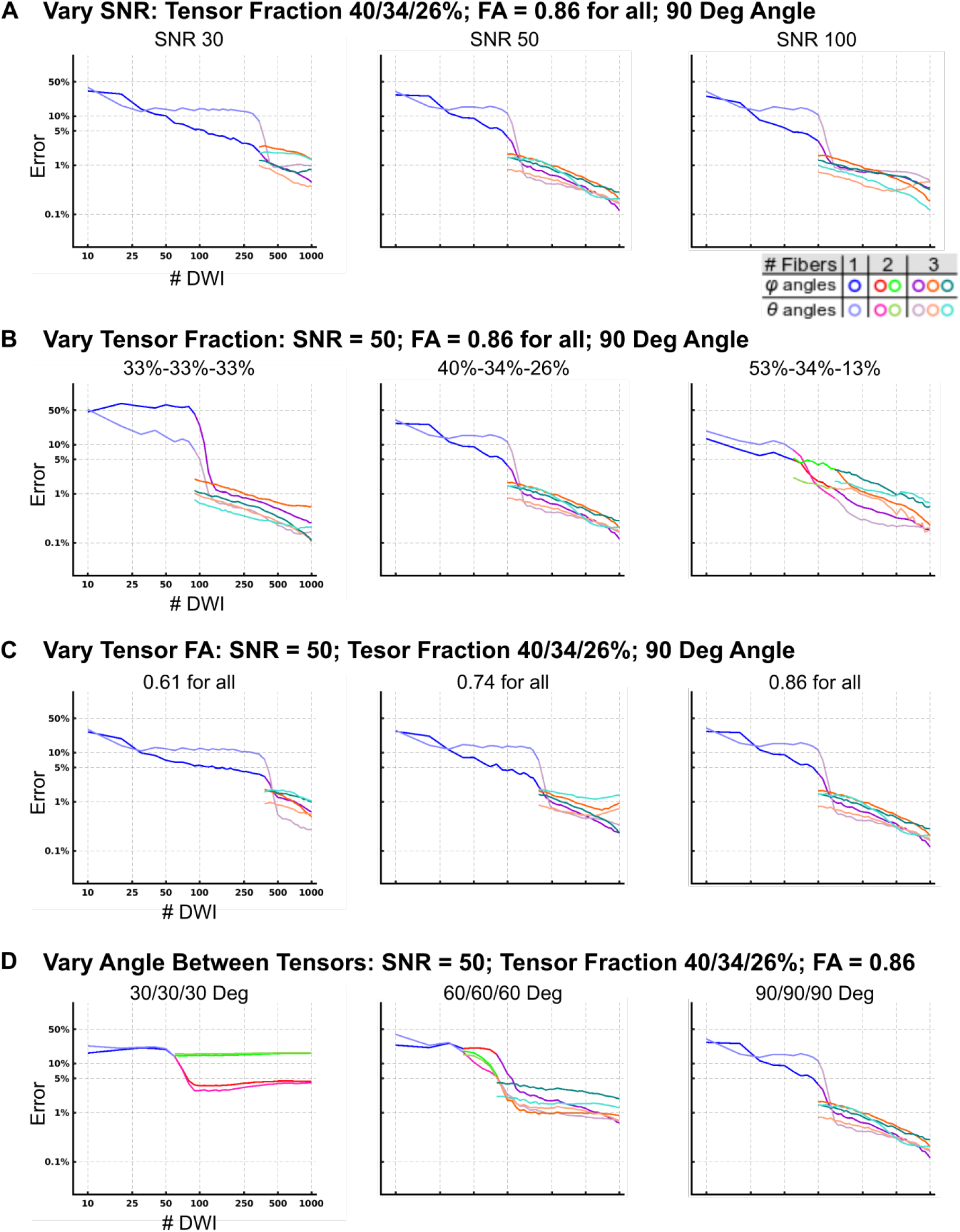
Accuracy of BaMM in variety of three tensor simulated data (A) Error estimation by BaMM for three crossing tensors at varying SNR. Mean error at each subsampling size was calculated, then plotted as the log of error. Plots are colored by the most frequent number of fibers estimated: subsamples with a single fiber direction are plotted in blue/sky blue, two fibers plotted red/pink and green/olive, 3 fibers are plotted in purple/lilac, orange/salmon, teal/cyan. (B) Vary tensor Fraction. (C) Vary tensor FA. (D) Vary angle between tensors.

**Figure 11:**
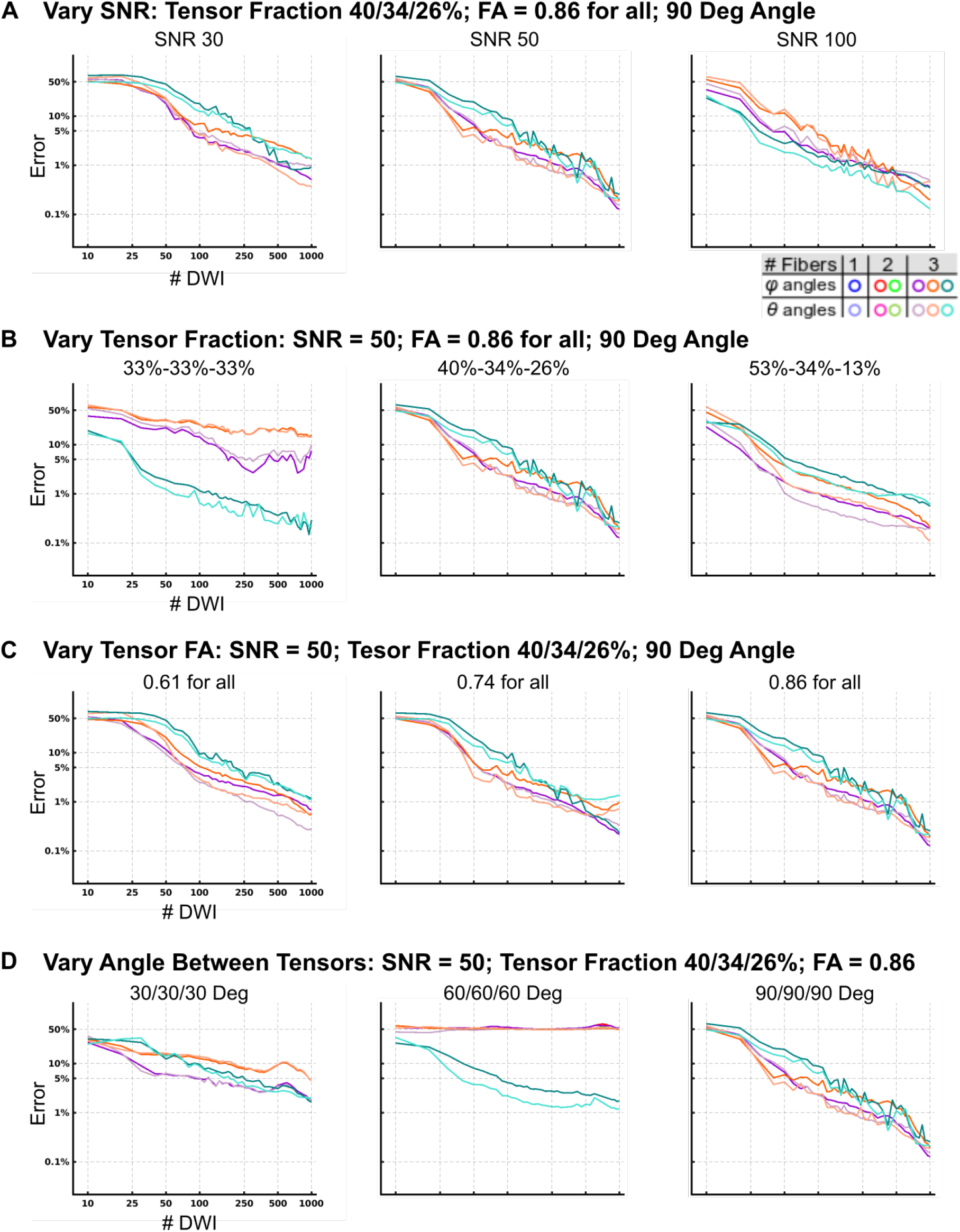
Accuracy of BPX in variety of three tensor simulated data (A) Error estimation by BPX for three crossing tensors at varying SNR. Mean error at each subsampling size was calculated, then plotted as the log of error. Plots are colored by the most frequent number of fibers estimated: subsamples with a single fiber direction are plotted in blue/sky blue, two fibers plotted red/pink and green/olive, 3 fibers are plotted in purple/lilac, orange/salmon, teal/cyan. (B) Vary tensor Fraction. (C) Vary tensor FA. (D) Vary angle between tensors.

**Figure 12:**
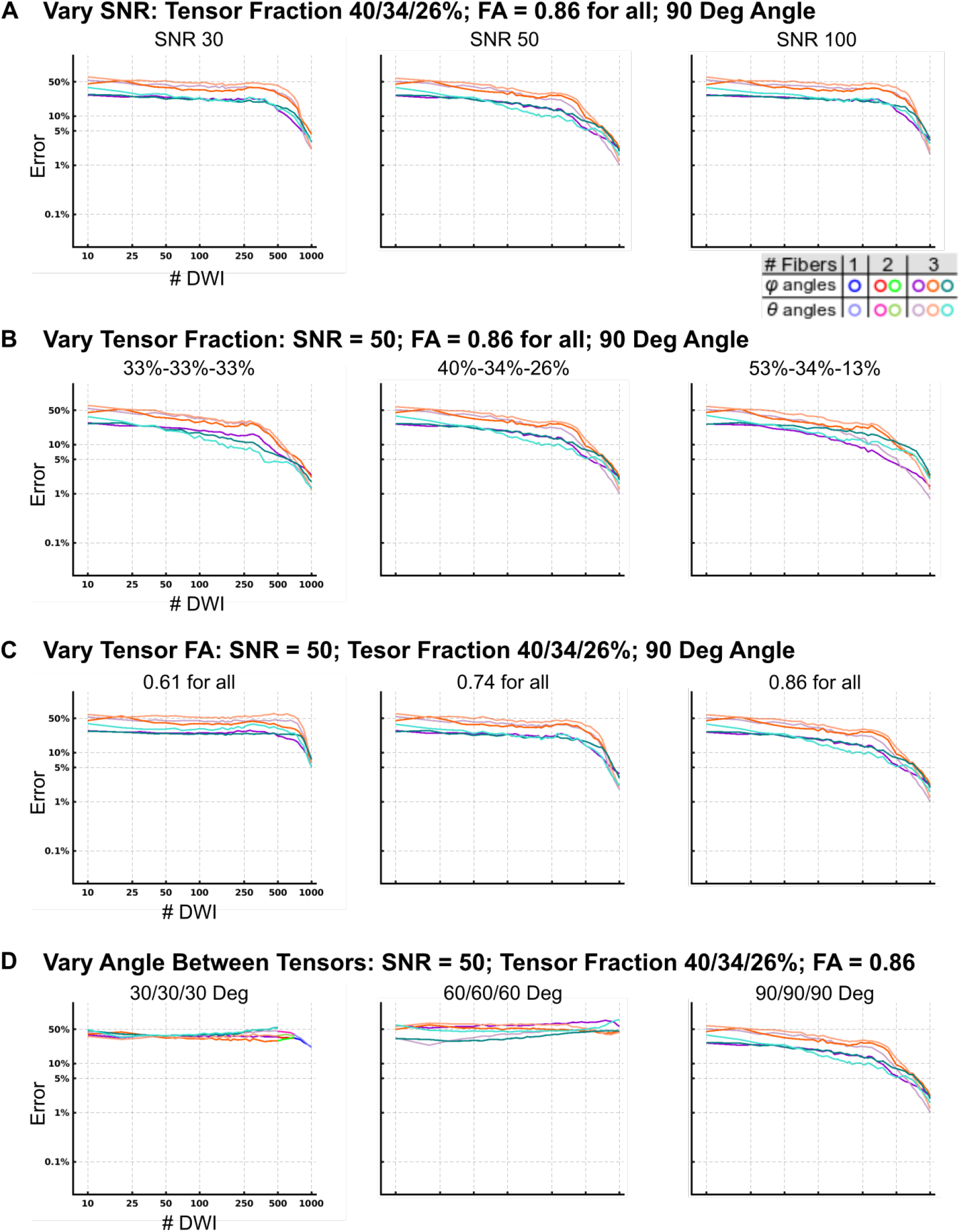
Accuracy of QBI in variety of three tensor simulated data (A) Error estimation by QBI for three crossing tensors at varying SNR. Mean error at each subsampling size was calculated, then plotted as the log of error. Plots are colored by the most frequent number of fibers estimated: subsamples with a single fiber direction are plotted in blue/sky blue, two fibers plotted red/pink and green/olive, 3 fibers are plotted in purple/lilac, orange/salmon, teal/cyan. (B) Vary tensor Fraction. (C) Vary tensor FA. (D) Vary angle between tensors.

**Figure 13:**
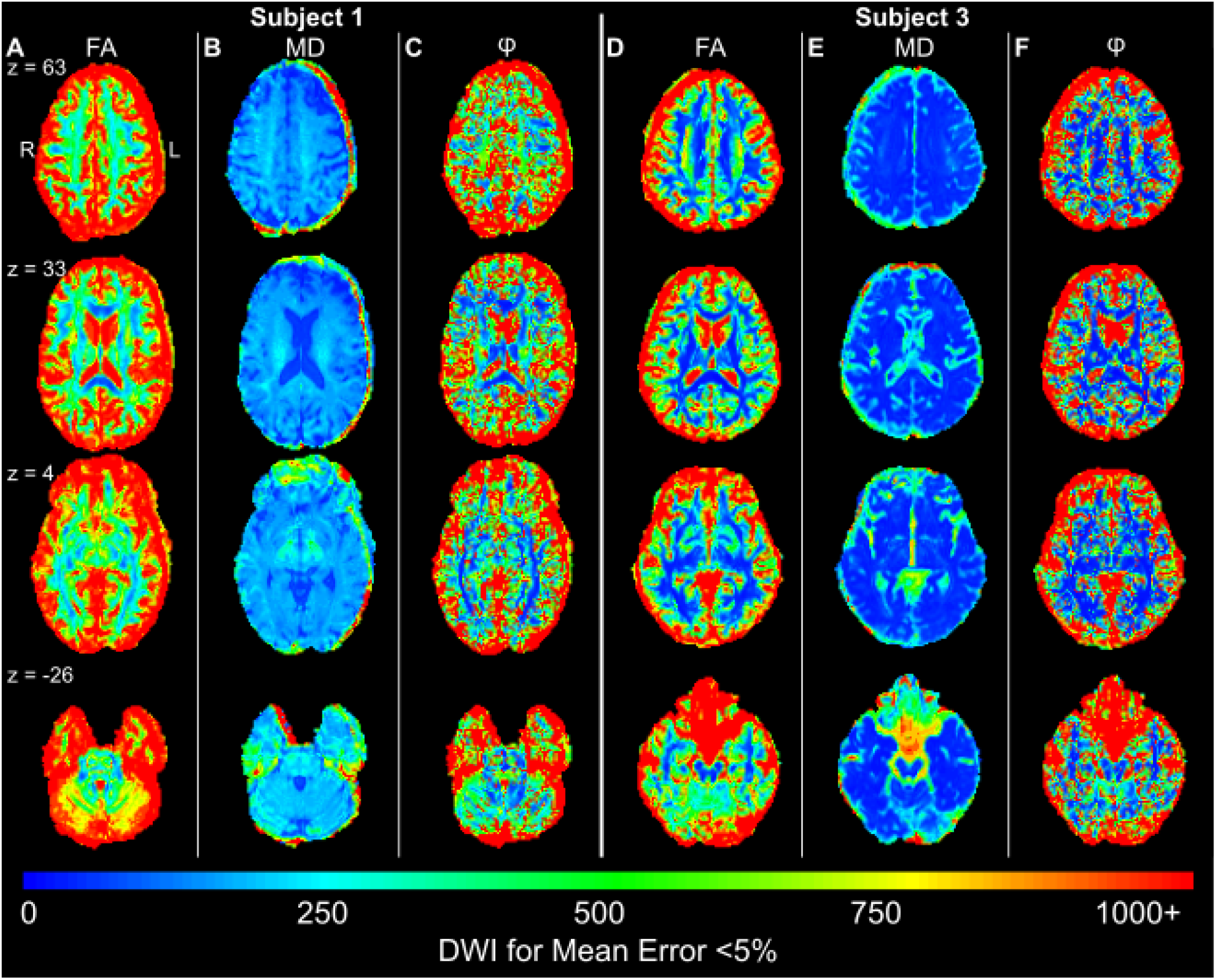
LLS whole-brain reliability map for Mean Error < 5%, Subject 1 & and Subject 3 (A) The color scale shows the number of DWI measurements needed to achieve a voxel-wise error less than 5% in FA. Error is calculated relative to the mean FA found using the entire sample. Results for (B) RD, (C) AD, (D) MD, and (E) angle *φ* are shown.

**Figure 14:**
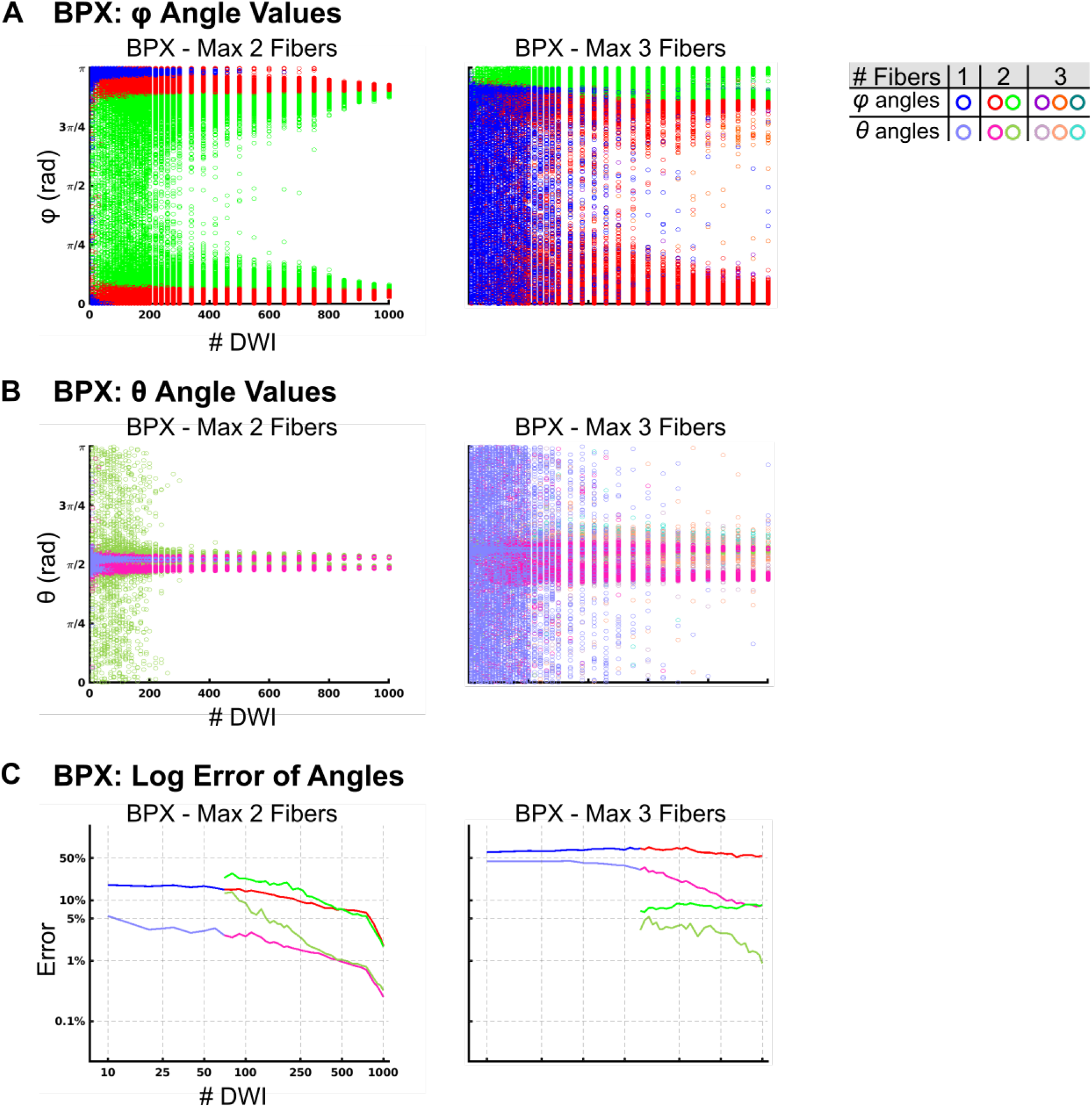
Reliability of BPX Subject 2 Corpus Callosum with max 3 fibers The same ROI as in figure 7 in subject 2 isanalyzed. (A) *φ* Angle estimations by BedpostX (BPX) with max two or three fibers. (B) *θ* Angle estimations by BPX with max two or three fibers. (C) Log Error estimation by BPX with two or three fibers. Error is calculated relative to the mean *φ* or *θ* found using the entire sample.

**Figure 15:**
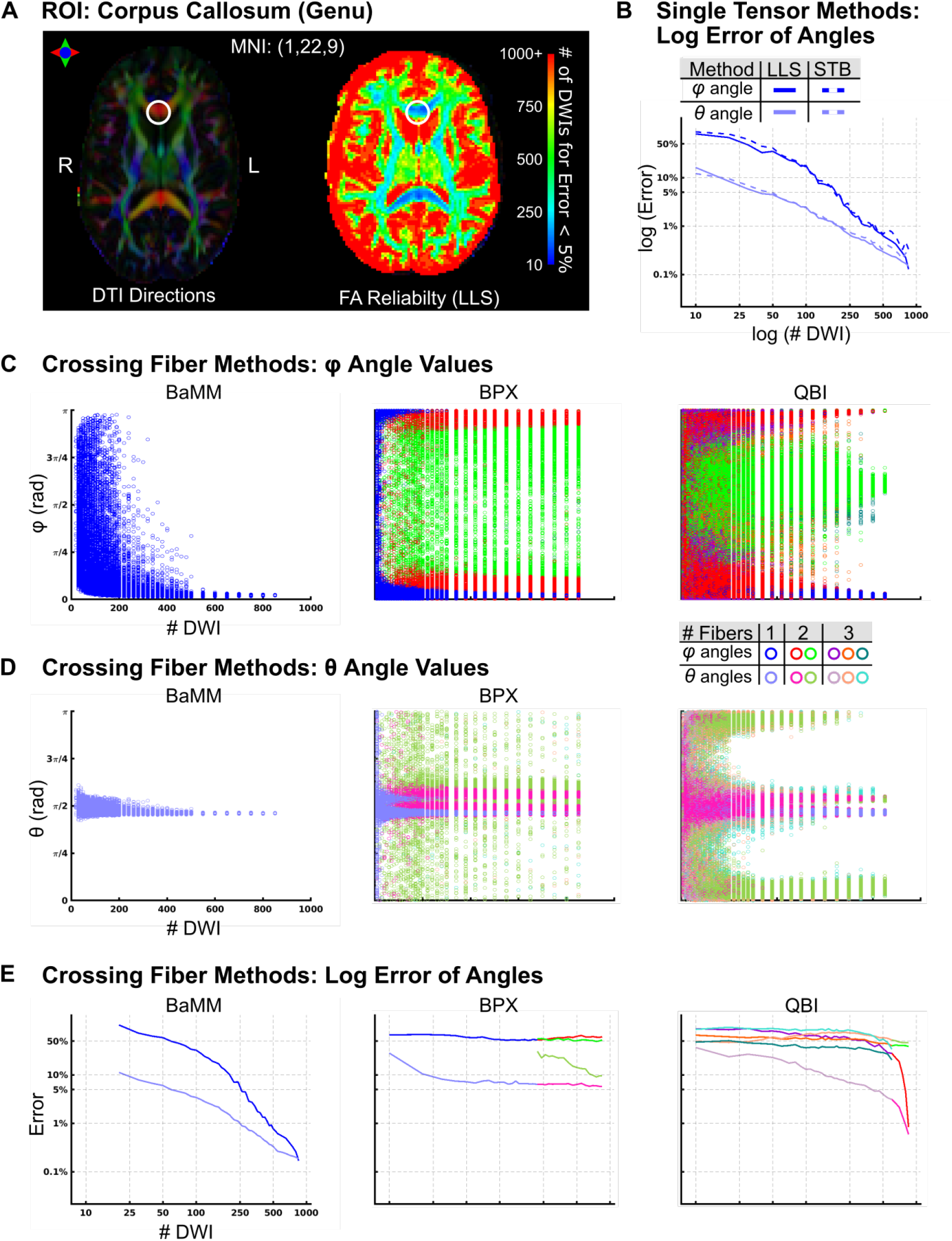
Reliability of Diffusion Measures in the Genu of the Corpus Callosum, Subject 1 (A)The locus of the analyzed voxel (MNI: 1, 22, 9) is marked with a circle. LLS FA reliability map as in Figure 6A. (B) Log Error of angle estimation by LLS and STB. (C) *φ* Angle estimations by Bayesian Multi-tensor Model-selection (BaMM), BedpostX (BPX), and ConstantSolid Angle Q-Ball Imaging (QBI). (D) *θ* Angle estimations by BaMM, BPX, and QBI. (E) Log Error estimation by BaMM, BPX, and QBI. Error is calculated relative to the mean *φ* or *θ* found using the entire sample

**Figure 16:**
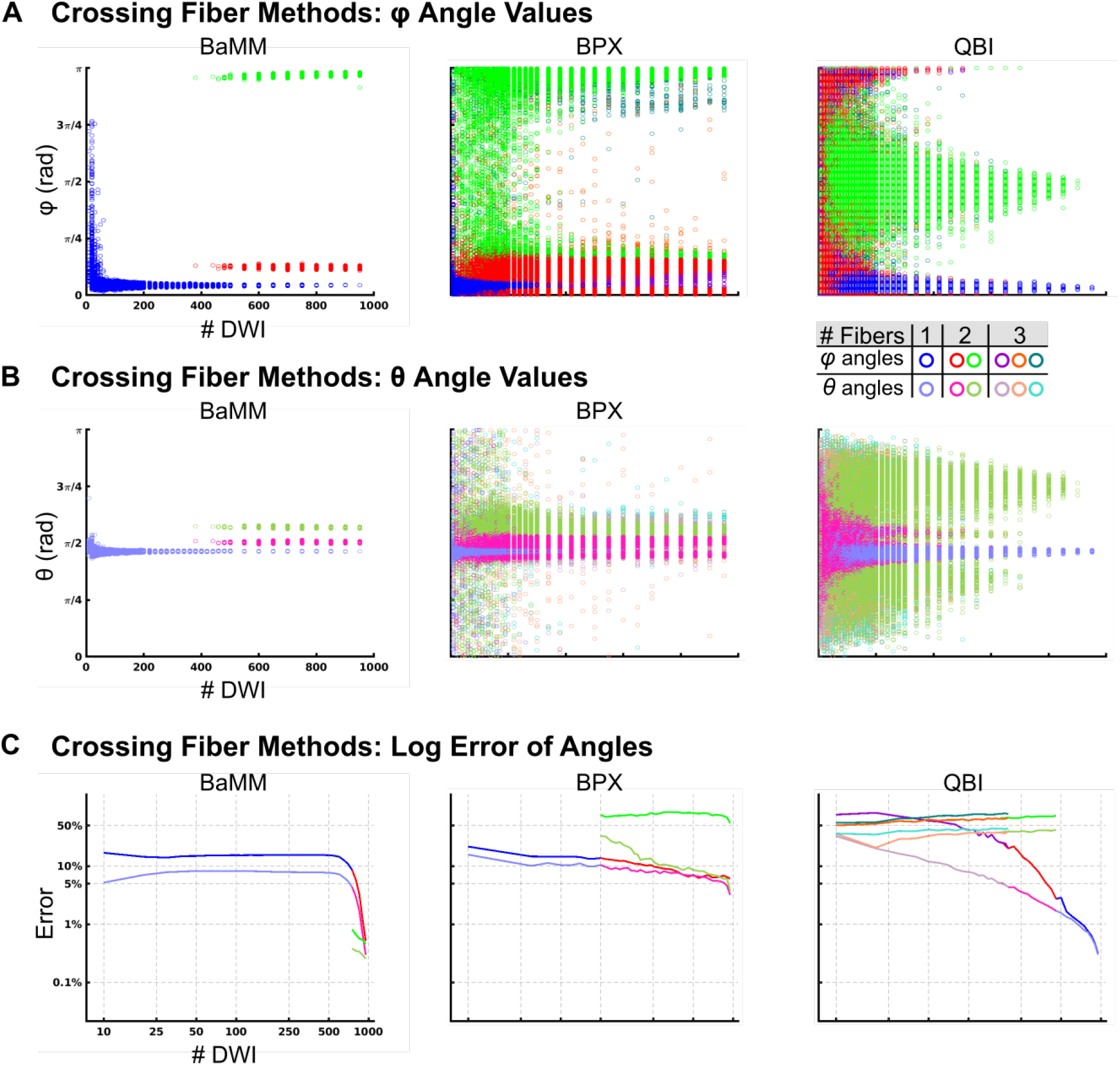
Reliability of Diffusion Measures in the Genu of the Corpus Callosum, Subject 1 Subject 1 was rescanned using a 1020 DWI sequence with 960 unique B-vector direction. The data was analyzed in native space, and the same ROI as in Figure S14 was tested. (A) *φ* Angle estimations by Bayesian Multi-tensor Model-selection (BaMM), BedpostX (BPX), and ConstantSolid Angle Q-Ball Imaging (QBI). (B) *θ* Angle estimations by BaMM, BPX, and QBI. (C) Log Error estimation by BaMM, BPX, and QBI. Error is calculated relative to the mean *φ* or *θ* found using the entire sample

**Figure 17:**
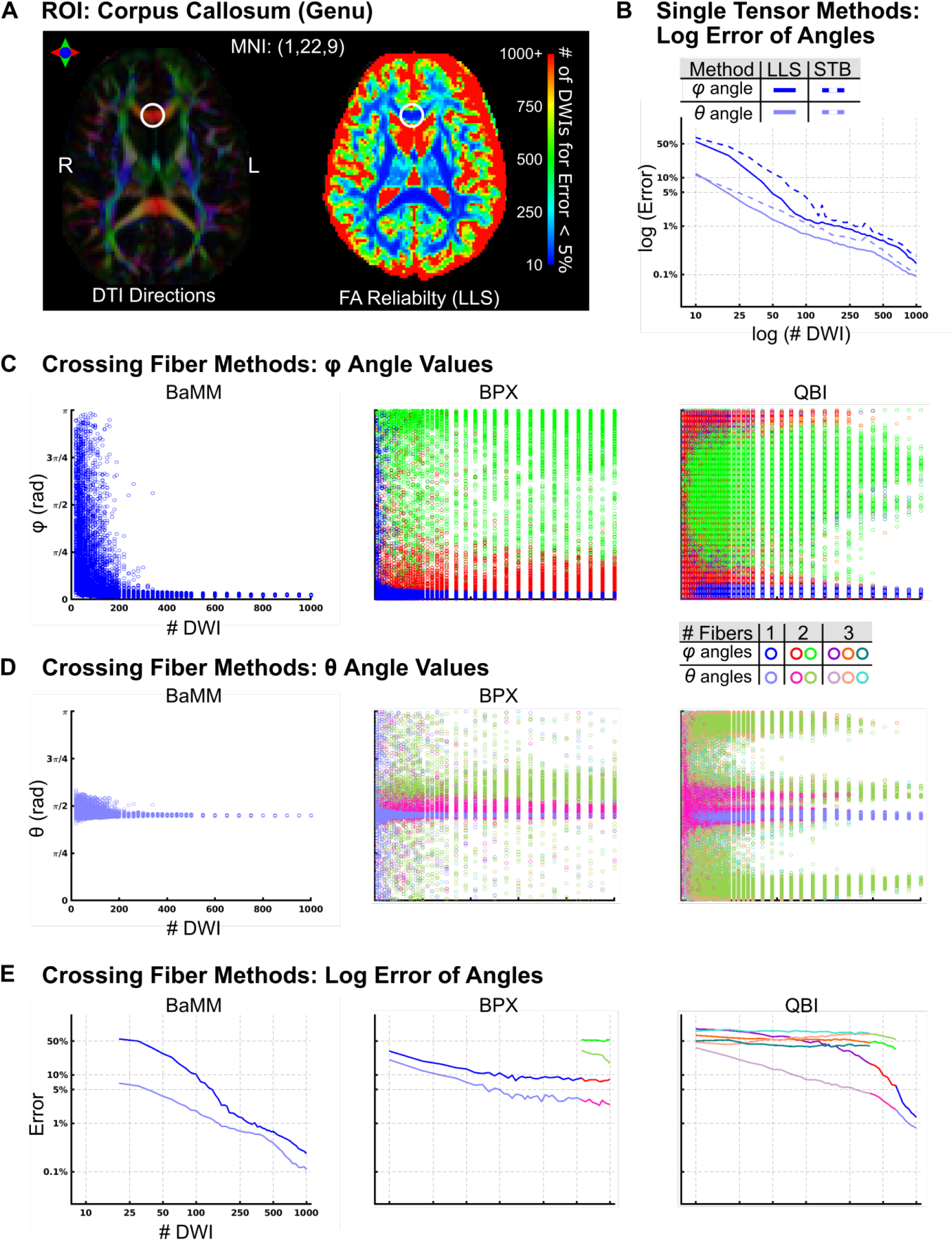
Reliability of Diffusion Measures in the Genu of the Corpus Callosum, Subject 3 (A)The locus of the analyzed voxel (MNI: 1, 22, 9) is marked with a circle. LLS FA reliability map as in Figure 6A. (B) Log Error of angle estimation by LLS and STB. (C) *φ* Angle estimations by Bayesian Multi-tensor Model-selection (BaMM), BedpostX (BPX), and ConstantSolid Angle Q-Ball Imaging (QBI). (D) *θ* Angle estimations by BaMM, BPX, and QBI. (E) Log Error estimation by BaMM, BPX, and QBI. Error is calculated relative to the mean φ or θ found using the entire sample

**Figure 18:**
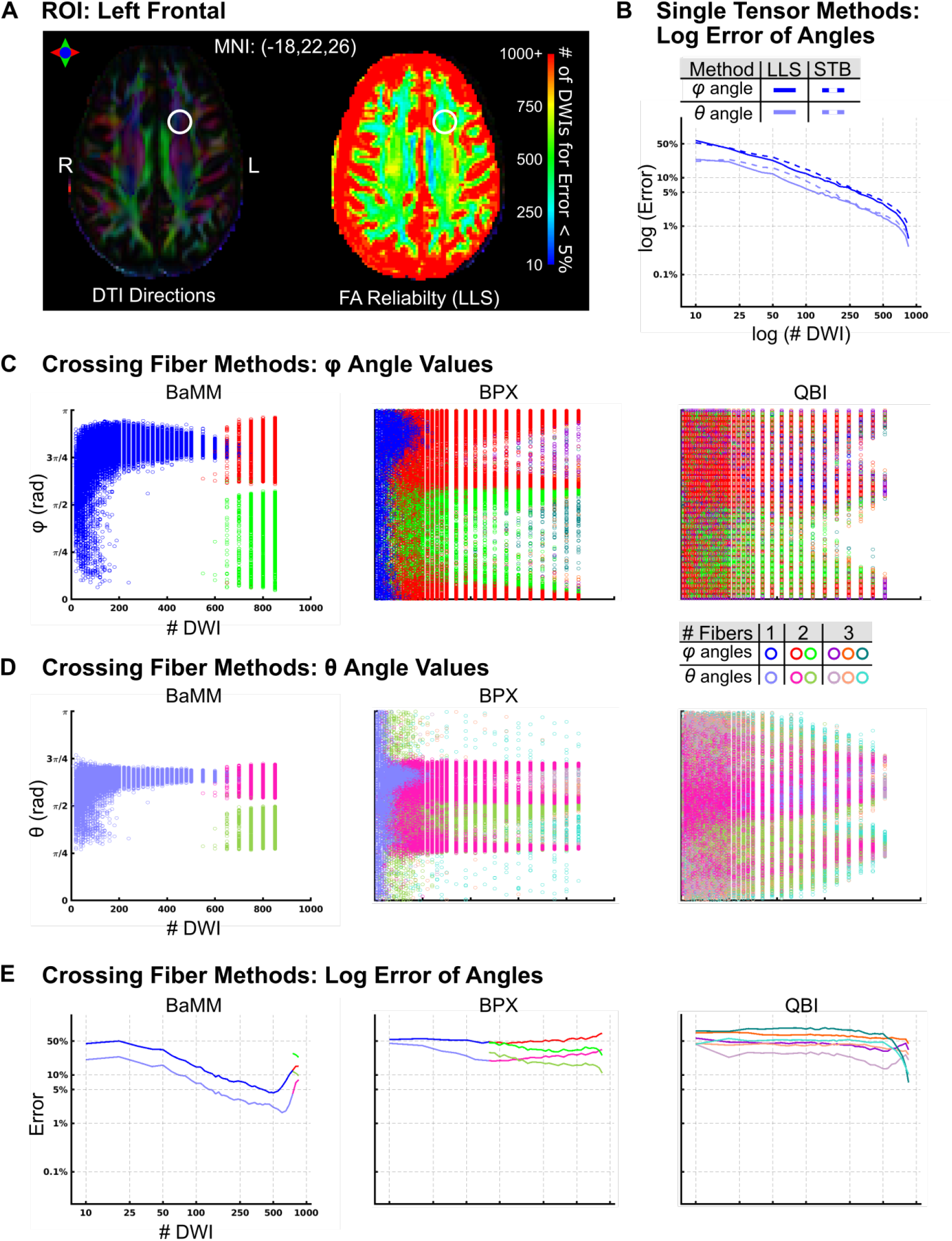
Reliability of Diffusion Measures in the Left Frontal White Matter, Subject 1 (A)The locus of the analyzed voxel (MNI: 18,22,26) is marked with a circle. LLS FA reliability map as in Figure 6A. (B) Log Error of angle estimation by LLS and STB. (C) *φ* Angle estimations by Bayesian Multi-tensor Model-selection (BaMM), BedpostX (BPX), and Constant Solid Angle Q-Ball Imaging (QBI). (D) *θ* Angle estimations by BaMM, BPX, and QBI. (E) Log Error estimation by BaMM, BPX, and QBI. Error is calculated relative to the mean *φ* or *θ* found using the entire sample

**Figure 19:**
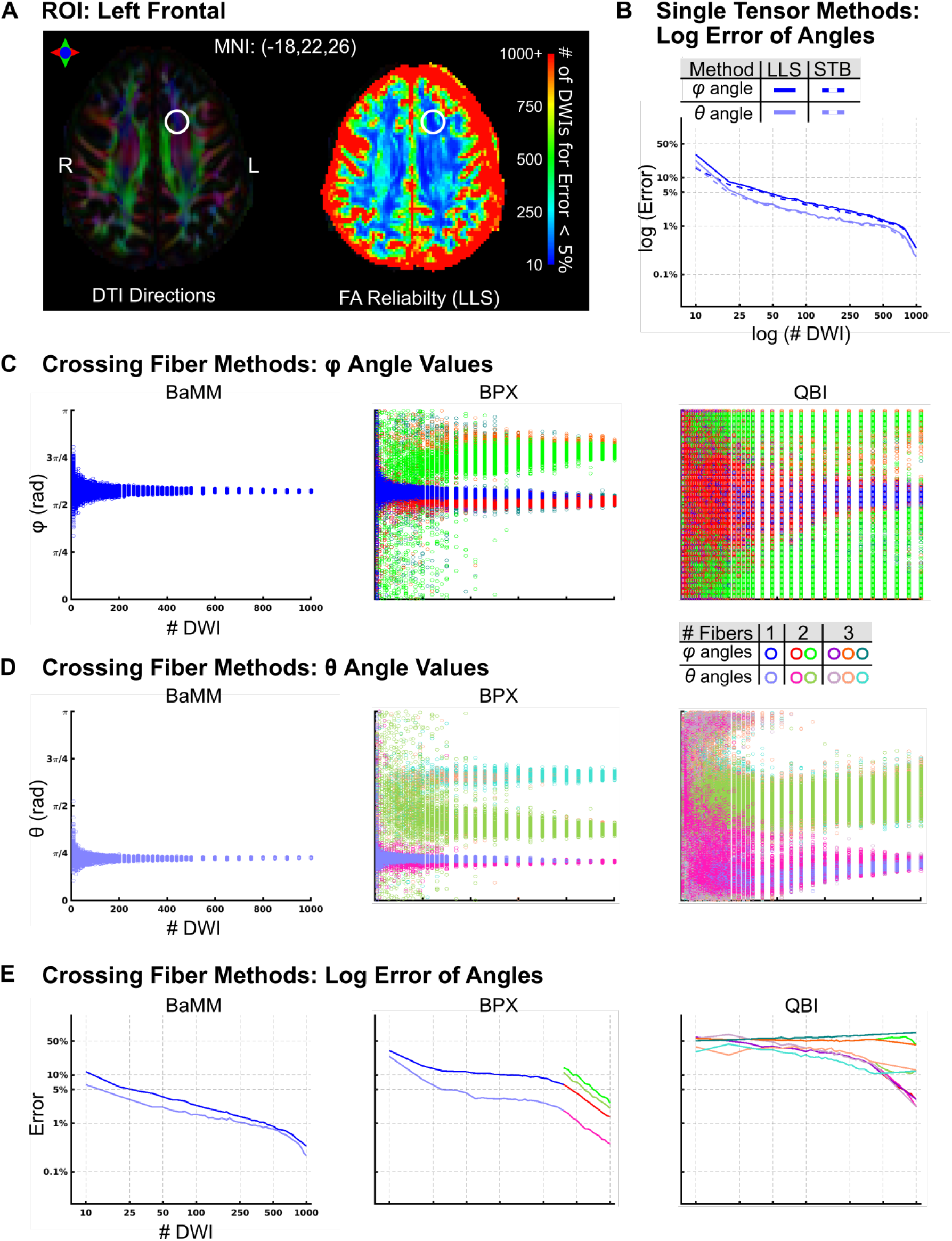
Reliability of Diffusion Measures in the Left Frontal White Matter, Subject 3 (A) The locus of the analyzed voxel (MNI: 18,22,26) is marked with a circle. LLS FA reliability map as in Figure 6A. (B) Log Error of angle estimation by LLS and STB. (C) *φ* Angle estimations by Bayesian Multi-tensor Model-selection (BaMM), BedpostX (BPX), and Constant Solid Angle Q-Ball Imaging (QBI). (D) *θ* Angle estimations by BaMM, BPX, and QBI. (E) Log Error estimation by BaMM, BPX, and QBI. Error is calculated relative to the mean *φ* or *θ* found using the entire sample

**Figure 20:**
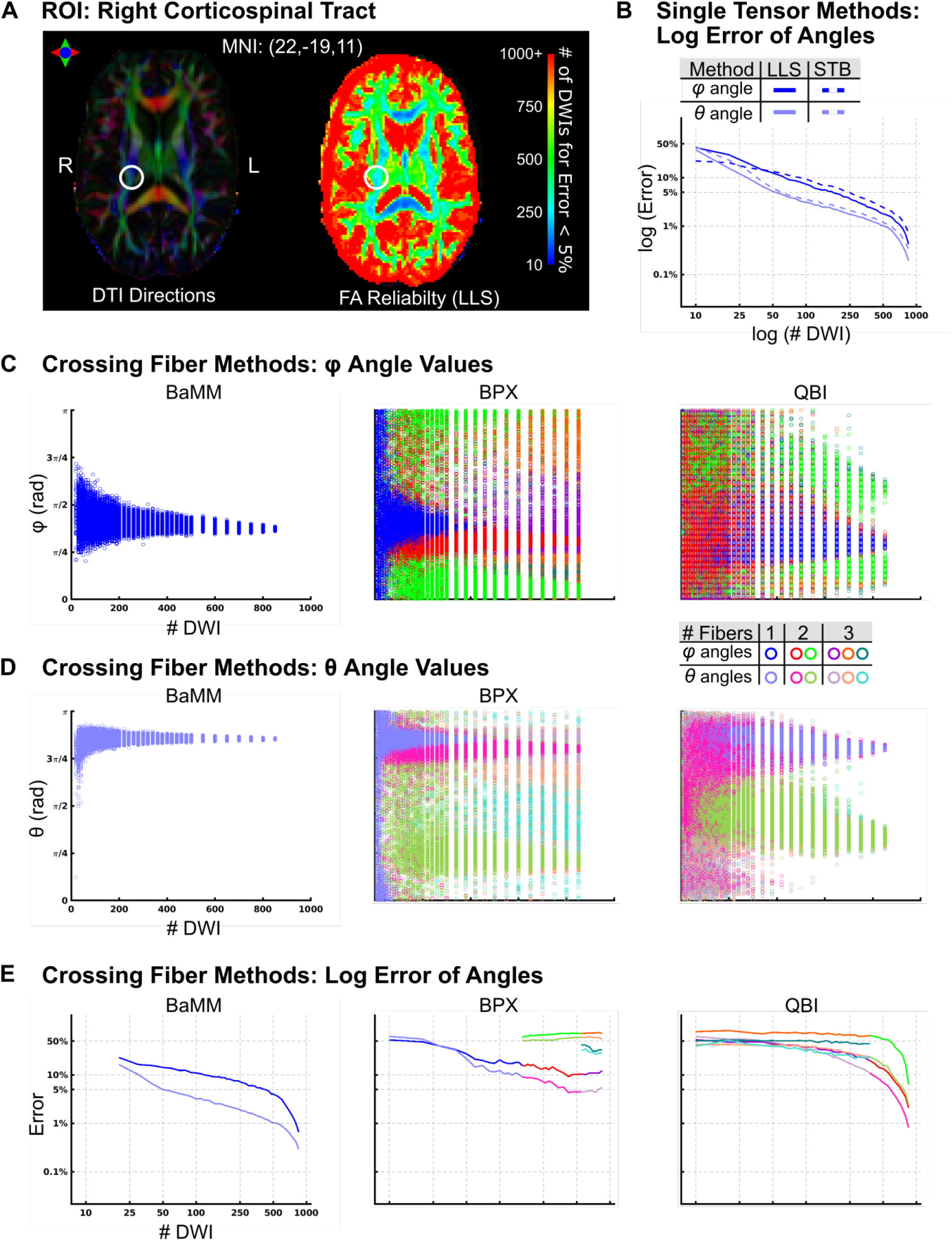
Reliability of Diffusion Measures in Right Corticospinal Tract, Subject 1 (A) The locus of the analyzed voxel (MNI: −22, −19,11) is marked with a circle. LLS FA reliability map as in Figure 6A. (B) Log Error of angle estimation by LLS and STB. (C) *φ* Angle estimations by Bayesian Multi-tensor Model-selection (BaMM), BedpostX (BPX), and ConstantSolid Angle Q-Ball Imaging (QBI). (D) *θ* Angle estimations by BaMM, BPX, and QBI. (E) Log Error estimation by BaMM, BPX, and QBI. Error is calculated relative to the mean *φ* or *θ* found using the entire sample.

**Figure 21:**
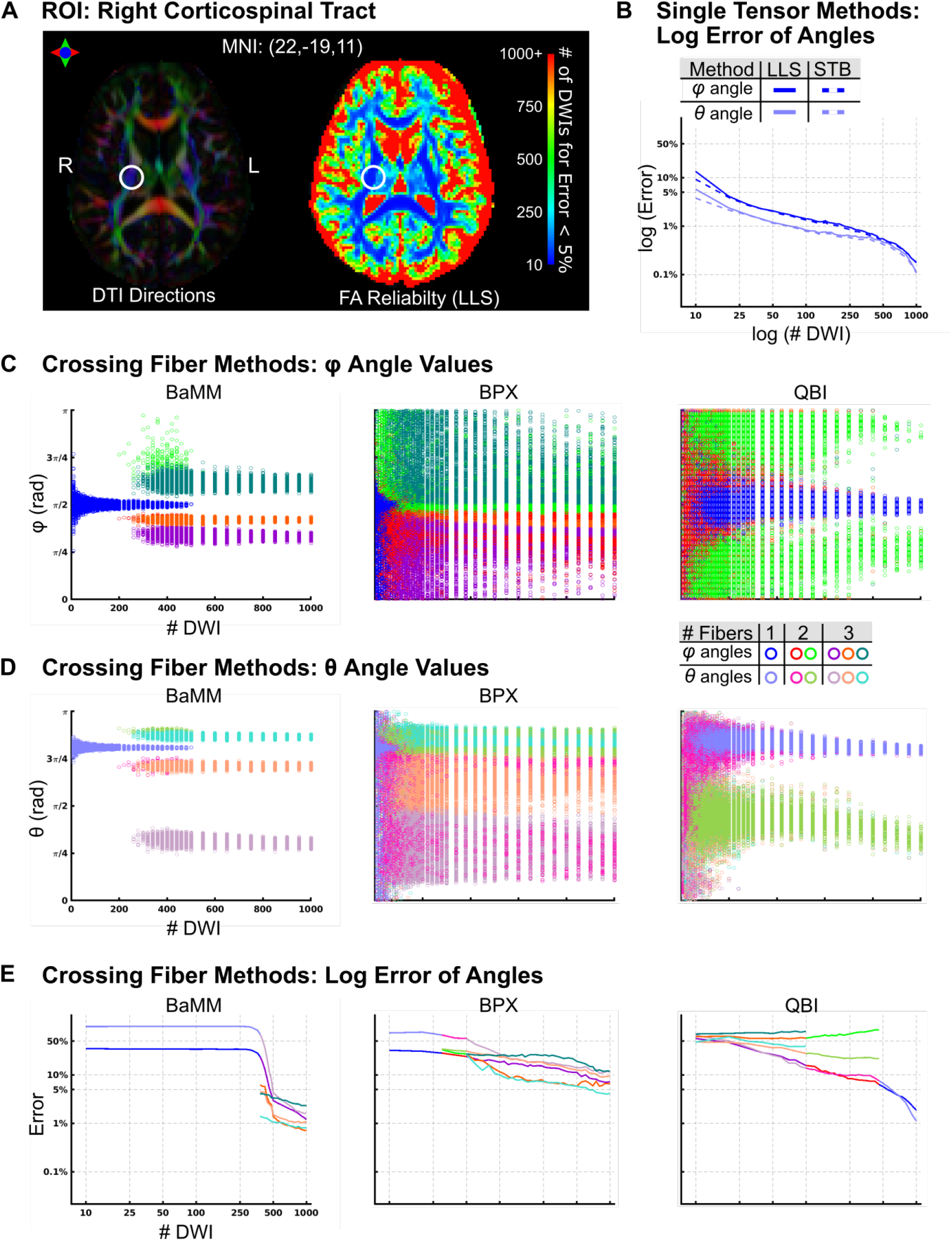
Reliability of Diffusion Measures in Right Corticospinal Tract, Subject 3 (A) The locus of the analyzed voxel (MNI: −22, −19,11) is marked with a circle. LLS FA reliability map as in Figure 6A. (B) Log Error of angle estimation by LLS and STB. (C) *φ* Angle estimations by Bayesian Multi-tensor Model-selection (BaMM), BedpostX (BPX), and ConstantSolid Angle Q-Ball Imaging (QBI). (D) *θ* Angle estimations by BaMM, BPX, and QBI. (E) Log Error estimation by BaMM, BPX, and QBI. Error is calculated relative to the mean *φ* or *θ* found using the entire sample.

